# The role of the synaptic vesicle protein SV2A in regulating mitochondrial morphology and autophagy

**DOI:** 10.1101/2024.07.30.605753

**Authors:** Marko Jörg, Jonas S. Reichert, Karin Pauly, Ute Distler, Stephan Tenzer, Odile Bartholomé, Bernard Rogister, Andreas Kern, Christian Behl, Martón Gellérie, Christoph Cremer, Sandra Ritz, Philipp Peslalz, Bernd Plietker, Kristina Friedland

**Affiliations:** Department of Pharmacology and Toxicology, Institute of Pharmaceutical and Biomedical Sciences, Johannes Gutenberg University Mainz, D-55128 Mainz; Core Facility for Mass Spectrometry, Institute of Immunology, University Medical Centre of the Johannes-Gutenberg University, Mainz, Germany; GIGA Neurosciences, University of Liege, CHU B36, 1 Avenue de L’hôpital, 4000, Liège, Belgium; Department of Neurology, CHU Liège, Liège, Belgium; Institute of Pathobiochemistry, University Medical Center of the Johannes Gutenberg University, Duesbergweg 6, 55128 Mainz, Germany; Institute of Molecular Biology gGmbH (IMB), Mainz, Germany; Organic Chemistry I, Faculty of Chemistry and Food Chemistry, TU Dresden, Bergstrasse 66, Dresden DE-01069, Germany

**Keywords:** SV2A, Mitochondria, Cell biology, Autophagy, cellular organelles

## Abstract

The synaptic vesicle glycoprotein 2A (SV2A) is a transmembrane protein of synaptic vesicles. It is involved in key functions of neurons, focused on the regulation of neurotransmitter release. Here we report three novel findings suggesting a completely new role of SV2A. First, we demonstrate that SV2A is localized at the outer mitochondrial membrane (OMM) using confocal and super-resolution microscopy. Second, Inactivation of SV2A in our cell and animal models leads to fragmented mitochondria. In addition, SV2A also affects the basal autophagic flux as well as mitophagy. Third, using proteomics analysis we demonstrate that SV2A interacts with the fission factor DRP1 and the autophagy factor ATG9A. Using AlphaFold3 we provide a first glimpse of the molecular interaction between DRP1 and SV2A. Our findings demonstrate that SV2A is not only a vesicular protein but also a mitochondrial protein in the OMM with defined functions regulating mitochondrial morphology and autophagy.

## Introduction

Mitochondria play a crucial role in energy metabolism. They generate ATP via oxidative phosphorylation (OXPHOS) and build a highly dynamic network^1^. Mitochondria are double membrane organelles composed of an outer mitochondrial membrane (OMM) which is separated from the inner mitochondrial membrane (IMM) by the intermembrane space (IMS)^2^. The IMM encloses the mitochondrial matrix and forms invaginations called cristae where the OXPHOS system is located. The functional plasticity of mitochondria is linked to their morphology. Fission and fusion events are important for the regulation of mitochondrial morphology ranging from tubular networks to small fragmented mitochondria^3,4^. These membrane-remodeling events are mediated by conserved large dynamin-like GTPase proteins such as mitofusin 1 and 2 (MFN1, MFN2) for OMM fusion and the optic atrophy protein 1 (OPA1) for IMM fusion orchestrating mitochondrial fusion ^5^. Dynamin-related protein (DRP1) is responsible for fission. Once recruited to the OMM by receptor proteins such as mitochondrial fission protein 1 (FIS1), mitochondrial fission factor (MFF) or mitochondrial dynamic proteins MID49/51^6–8^, DRP1 forms helical oligomers and assembles into polymers which encircle mitochondria channeling energy from GTP binding, hydrolysis and nucleotide exchange into mechanochemical constriction resulting in mitochondrial fission. Recently, several cryo EM structures provide detailed understanding about the structure of DRP1 itself as monomer, oligomer or polymer as well as its interaction with the respective receptors^9–11^. DRP1 consists of a GTPase or G domain, where GTP binding and hydrolysis occurs and the stalk, comprised of the middle domain (MD) and GTPase effector domain (GED) connected by the bundle signaling element (BSE)^9–12^. In addition, DRP1 has a unique variable domain composed of 136 intrinsically disordered residues. The conformational change of the G domain due to GTP binding stabilizes DRP1 oligomeric structure at the OMM. MD is involved in self-assembly of DRP1 into dimers, tetramers and higher order oligomers^9-12^. DRP1-DRP1 dimerization is controlled by three helices of the MD and 1 helices of the GED^11^. Kalia et al. identified binding sites of MID49 in the G domain as well as the MD^9^.

Mitochondrial fragmentation is often linked to mitochondrial dysfunction^5^. However, mitochondrial fission is also required for mitochondrial motility, quality control and mtDNA inheritance ^5^. A fused mitochondrial network is considered to allow matrix component distribution, stimulation of OXPHOS activity resulting in enhanced ATP levels and is mainly associated with cell survival mechanisms^2,4,13^. Damaged and fragmented mitochondria are removed by one form of autophagy, mitophagy. Autophagy is able to selectively eliminate unwanted, potentially harmful cytosolic material, such as damaged mitochondria or protein aggregates ^14^. The whole process of autophagy can be divided into 4 different stages: autophagy initiation including phagophore elongation, autophagosome formation, autophagosome maturation and lysosomal degradation^15^. Induction of autophagy results in the recruitment of autophagy-related proteins (ATGs) resulting in the assembly of the phagophore ^14^. ATG9 is of special interest since it is the only conserved transmembrane protein mainly localized in Golgi-derived vesicles functioning as the initial membrane source for autophagosome initiation^16,17^. It is also important for mitophagy initiation^18^.

An integral constituent of vertebrate synaptic vesicle membranes is the synaptic vesicle protein 2A (SV2A) which belongs to the Synaptic Vesicle protein 2 family (SV2)^19–21^. SV2A is considered to play a role in calcium-dependent exocytosis, endocytosis, neurotransmitter loading/retention in synaptic vesicles, and synaptic vesicle priming as well as transport of vesicle constituents^20–23^. SV2A harbors 12 transmembrane helices^24^. In addition to the transmembrane domain (TMD) core and a cytosolic N-terminal region, a large fourth luminal domain (LD4) are present. Two cytoplasmic horizontal helices, H1 and H3, constitute the cytoplasmic domain. The long loop extending from H3 continues to the horizontal H4 of the C-terminal half^24^. The junction between the N- and C-halves is disordered in the cryo-EM map^24^. SV2A was shown to form protein-protein interactions with synaptic vesicle proteins such as synaptophysin, synaptobrevin 2 and synaptotagmin 1^22,25^. SV2A interacts with synaptotagmin 1 via its cytoplasmatic N-terminal region^22^.

The importance of SV2A for proper central nervous system (CNS) functioning is demonstrated by severe defects in a genome-wide SV2A knockout (KO) mouse model^26^. 50% of these animals die during their first postnatal days. The remaining pups die within three weeks expressing severe seizures ^20,26^. The role of SV2A for epilepsy is further supported by the clinical highly effective anticonvulsive drugs levetiracetam and brivaracetam, both SV2A ligands^27–29^. Levetiracetam (LEV) is a well-tolerated drug recommended by clinical guidelines for different forms of epilepsy in adults and children^30^. In Alzheimer’s disease (AD), levetiracetam is discussed to improve hippocampal hyperexcitability occurring early in the disease process^31,32^. These effects are mainly attributed to changes in neurotransmitter release via modulation of the vesicular function of SV2A. However, recent data from our group demonstrates that LEV also improved mitochondrial function in a neuronal AD cell model leading to elongated mitochondria, reduced opening of the mitochondrial transition pore (MPTP) and rise in ATP levels^33^. Furthermore, siRNA-mediated SV2A knockdown in neuronal cells leads to mitochondrial deficits such as impaired mitochondrial morphology, e.g. mitochondrial fragmentation as well as reduced ATP levels. Furthermore, we demonstrated using highly purified mitochondria from mouse brain that SV2A might be localized in mitochondrial membranes. Given the dynamic nature of mitochondria involving continuous membrane fission and fusion akin to synaptic vesicles, we hypothesize that SV2A may exert similar regulatory effects on mitochondrial dynamics. Recent findings indicated mitochondrial localization of a subset of SNARE proteins which are well recognized as regulators of presynaptic neurotransmitter release and endocytotic recycling of synaptic vesicles^34^. Syntaxin17 (SYN17) knockdown leads to elongated mitochondria affecting DRP1 localization and activation^35^. Syntaxin 4 (Stx4) was found to be localized on or in proximity to the mitochondrial surface suppressing mitochondrial fission^36^. Other SNARE proteins known for their mitochondrial localization are e.g. SNAP23 as well VAMPB1^34^. They are also known to affect / modulate / influence autophagy. Furthermore, the mitochondrial fission factor DRP1 was found in conjunction with Bcl-XL, to influence synaptic vesicles, modulating endocytosis, vesicular membrane dynamics, and presynaptic plasticity in the hippocampus^37,38^. Therefore, we hypothesized that SV2A might share some functions with the so called mitoSNAREs.

Here we test three major hypotheses using a broad set of methods ranging from confocal and super resolution microscopy to proteomics and modelling protein-protein interactions. First, since SV2A is a membrane protein, we speculated that SV2A might be localized at OMM. Second, we asked whether SV2A impacts mitochondrial morphology as well as autophagy. And third, we questioned if SV2A not only interacts with vesicular proteins but also with mitochondrial proteins or proteins involved in autophagy/mitophagy.

## Results

### SV2A is localized at OMM

To test our first hypothesis and to investigate the precise localization of SV2A within mitochondria and its distribution across mitochondrial compartments, we employed four different experimental approaches: newly developed prediction-based tools, confocal microscopy, super-resolution microscopy GSIDM, and isolated mitochondria subjected to increasing concentrations of the detergent digitonin, causing their disintegration into individual fractions. First, we used newly developed tools like iMLP and DeepLoc2.0 to predict the subcellular localization of SV2A. The iMLP prediction shows the presence of internal mitochondrial targeting signals in SV2A and its isoform (Supplementary Fig. 1). The DeepLoc2 algorithm predicted SV2A as a transmembrane domain protein, also locating it at the mitochondria (Supplementary Fig. 2). These initial predicted results prompt us to further elaborate on the precise localization of SV2A within the mitochondria.

For confocal microscopy, SV2A was immunostained with a primary monoclonal antibody against SV2A and a secondary antibody coupled with Alexa488, while mitochondria were labeled with MitoTracker Deep Red FM in SH-SY5Y cells. We observed a colocalization of SV2A with mitochondria, supported by the calculation of the Pearson’s correlation coefficient (Pearson’s r) of 0.75. A Pearson’s r value of ≥ 0.5 is indicative of colocalization between two stained proteins (Fig. 1A).

**Figure 1.**
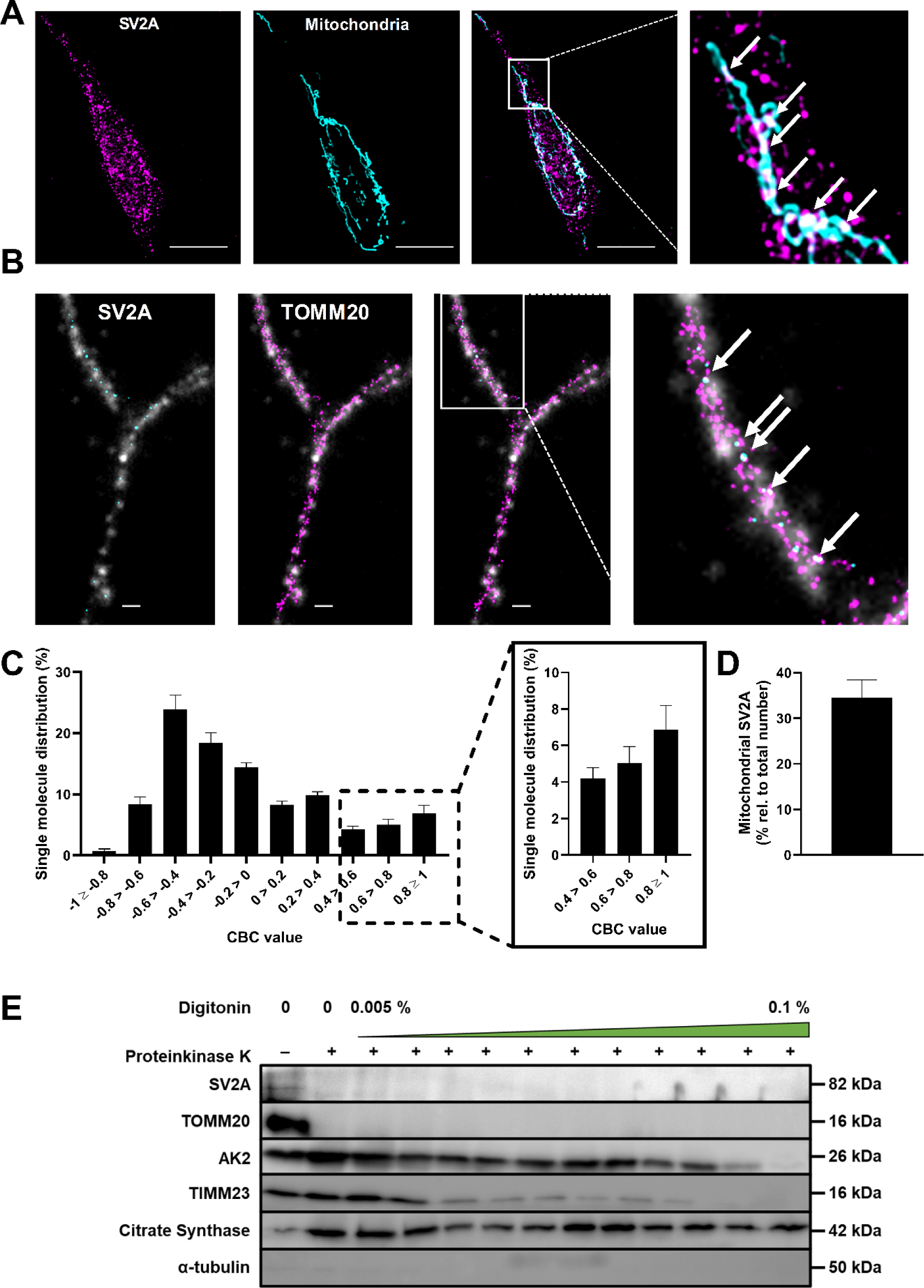
SV2A is localized at the OMM. A Representative confocal fluorescence images of double-immunolabeled human SH-SY5Y cells stained for SV2A and mitochondria using MitoTracker™ Deep Red FM. Colocalization of SV2A and mitochondria is indicated by white spots in the merged image (arrows). Scale bar: 10 µm. B Representative super-resolution images of human SH-SY5Y cells stained for SV2A and TOMM20 or TOMM20 and SV2A proteins with CBC values ≥0.5. A CBC value ≥0.5 indicates that these SV2A molecules colocalize with TOMM20. Mitochondrial area is displayed by a gray widefield image of TOMM20. Scale bar: 1 µm. C Histogram of CBC value distribution for the colocalization of SV2A and TOMM20. Enlarged are CBC values corresponding to the colocalization of the two proteins of interest. D Amount of SV2A molecules within mitochondrial areas compared to total number. E Representative Western blot analysis of mitochondrial fractions. For each mitochondrial fraction, a marker protein was detected to visualize the existing fractions per lane. SV2A (protein of interest), TOMM20 (OMM), AK2 (IMS), TIMM23 (IMM), Citrate synthase (matrix), α-tubulin (cytosolic marker). Data information: In (A-B) n= 11 ± SEM from 3 independent biological experiments. In (D) n = 11 ± SEM cells from 3 independent biological experiments. In (E) n= 3 independent biological experiments were analyzed.

To further corroborate these findings, we employed super-resolution GSDIM microscopy, which provides a resolution approximately 15 times higher in the xy direction than conventional confocal microscopy^39,40^. GSDIM microscopy was utilized to ascertain the precise localization of SV2A molecules within mitochondria and to assess the reliability of the recent confocal microscopy data. SV2A was again labeled using a monoclonal primary antibody and an ATTO488 coupled secondary antibody, while mitochondria were visualized using a primary antibody against the translocase of the outer mitochondrial membrane subunit 20 (TOMM20) and an Alexa647 coupled secondary antibody. Again, we observed colocalization of SV2A and TOMM20 in super-resolution images, which supports the findings obtained with confocal microscopy. The colocalization of SV2A and TOMM20 suggests that the two proteins are less than 40 nm apart (Fig. 1B, Supplementary Fig. 3). Considering that mitochondria are larger than 40 nm, our results strongly support the localization of SV2A within or at mitochondria.

To obtain more precise data regarding the number of SV2A molecules colocalized with TOMM20 and to determine whether SV2A is indeed a protein of the outer mitochondrial membrane (OMM), we calculated the coordination-based colocalization (CBC) for each coordinate of SV2A and TOMM20 using FIJI image software (Fig. 1C). The CBC, similar to the Pearson’s r, range from -1 to 1, with -1 representing anti-correlation and 1 representing colocalization. The closer the proteins are to each other, the higher the CBC value. The CBC histogram of SV2A and TOMM20 in control cells revealed that 14.1% shared CBC values ≥ 0.5, suggesting that SV2A might indeed be localized at the outer mitochondrial membrane (Fig. 1C,D).

To further validate these findings, we isolated mitochondria and exposed them to increasing concentrations of the detergent digitonin. At low concentrations, digitonin selectively permeabilizes the OMM, while at higher concentrations, it permeabilizes the inner membrane. The samples were then analyzed by Western blotting together with specific marker proteins for each mitochondrial compartment to identify SV2A in the respective mitochondrial fractions. Since SV2A reacts poorly to temperature (Supplementary Fig. 4), the SV2A samples were only heated to 70°C. As suspected, SV2A was found in the OMM, as confirmed by the presence of the SV2A band in the fraction containing the OMM marker TOMM20. Furthermore, SV2A disappeared along with the OMM marker TOMM20 as digitonin concentrations increased (Fig. 1E).

### SV2A knockdown leads to mitochondrial fragmentation in cell and animal model

To test the second hypothesis if SV2A might regulate mitochondrial morphology, we first employed a cell-based approach by knocking down SV2A using target-based siRNA. Control cells treated with scrambled siRNA serving as negative control demonstrated tubular mitochondrial networks associated with a low number of fragmented mitochondria, whereas mitochondria in SV2A KD cells appeared smaller, more fragmented, and swollen (Fig. 2A). In terms of mitochondrial length, SV2A KD exhibited a significant shift toward punctuated mitochondria and a decrease in tubular mitochondria, suggesting that it might play a crucial role in mitochondrial morphology. (Fig. 2A, Supplementary Fig. 5). Positive SV2A KD was validated using quantitative RT-PCR (qPCR) (Fig. 2B).

**Figure 2.**
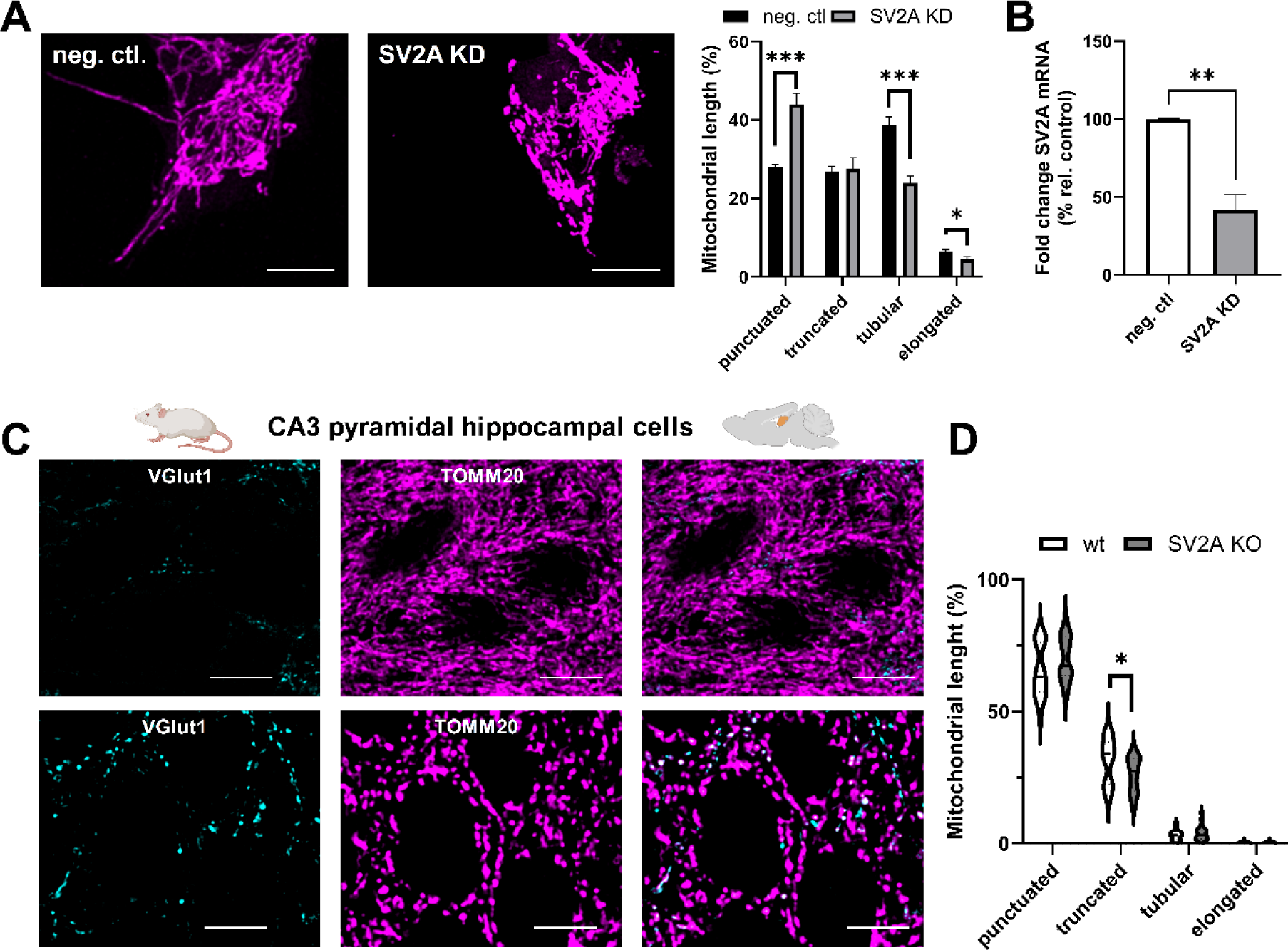
Loss of SV2A results in mitochondrial fragmentation and altered mitochondrial dynamics in SH-SY5Y cells and KO animals. A Representative confocal images of mitochondria in siRNA-induced SV2A KD cells and cells treated with scrambled siRNA (10 nM/48 h). Scale bar: 10 µM. B Length distribution of mitochondria in SV2A KD cells compared to cells treated with scrambled siRNA. C Analysis of SV2A mRNA levels in siRNA-induced SV2A knockdown cells (SH-SY5Y) confirmed knockdown of protein of interest using customized arrays for RT-qPCR. D Representative confocal fluorescence images of CA3 pyramidal cells of wildtype and CA3-specific SV2A KO male and female mice stained for VGlut1 and TOMM20. Scale bar: 10 µM. E Length distribution of mitochondria in CA3 pyramidal cells of wildtype and CA3-specific SV2A KO mice stained for VGlut1 and TOMM20. Scale bar: 10 µM. Data information: In (A-C) n = 6. In (E) 5 wt female and 5 wt male mice vs. 4 SV2A KO female and 4 SV2A male mice were analyzed. 3 slices / animal were analyzed. In (B,C and D) Data is expressed as mean ± SEM; student’s unpaired t-test (*p <0.01).

To further strengthen this hypothesis, we used a SV2A knockout (KO) mouse model. Given that homozygous SV2A KO results in the death of juvenile mice associated with seizures, we opted for the SV2A KO mouse model introduced by Menten-Dedoyart et al., which is unique in its hippocampus-restricted SV2A KO specifically targeting dentate gyrus and CA3 neurons^41^. The advantages of this mouse model over other SV2A KO mouse models are two-fold. First, mice of any age can be studied since these animals do not die prematurely. Second, these animals do not experience seizures, which would strongly affect mitochondrial morphology. Glutamatergic neurons of the hippocampal CA3 region were examined double-blinded using a confocal microscope to identify differences in mitochondrial morphology in the brains of 12-week-old SV2A KO and control animals. Glutamatergic neurons were visualized TOMM20 immunostaining with Alexa647 antibody. In control mice, mitochondria exhibited a tubular interconnected network, whereas mitochondria in SV2A KO mice were highly fragmented due to the loss of SV2A (Fig. 2C,D). Double-blinded measurements of mitochondrial length in SV2A KO hippocampi similarly reflected a shift in mitochondrial length towards smaller, punctuated mitochondria compared to respective controls. The percentage of punctuated mitochondria increased, accompanied by a substantial loss of mitochondria within the range of truncated mitochondria (Fig. 2D). The numbers of mitochondria classified as elongated and tubular were similar both SV2A KO and control mice.

### SV2A ligand LEV changes mitochondrial morphology and regulates a specific function of SV2A

Next, we investigated the effects of LEV (Fig. 3K), the SV2A ligand, on mitochondrial morphology. Since therapeutic plasma levels of LEV in mammals range from 23 µM to 176 µM and the maximum plasma concentration is reached after 1.3 hours, we chose to treat SH-SY5Y cells with 200 µM LEV for 2 h. LEV treatment resulted in a decrease in punctuated mitochondria and an increase in tubular mitochondria. The percentage of small punctuated mitochondria decreased by 7.4% during LEV treatment, while the percentages of tubular (+5.5%) and elongated (+1.7%) mitochondria increased (Fig. 3A,B). To investigate if LEV requires SV2A to alter mitochondrial morphology, we used confocal microscopy and treated our SV2A KD cells with LEV. The previously described effects, regarding mitochondrial elongation under LEV treatment, were completely abolished in SV2A KD cells (Fig. 3C,D).

**Figure 3.**
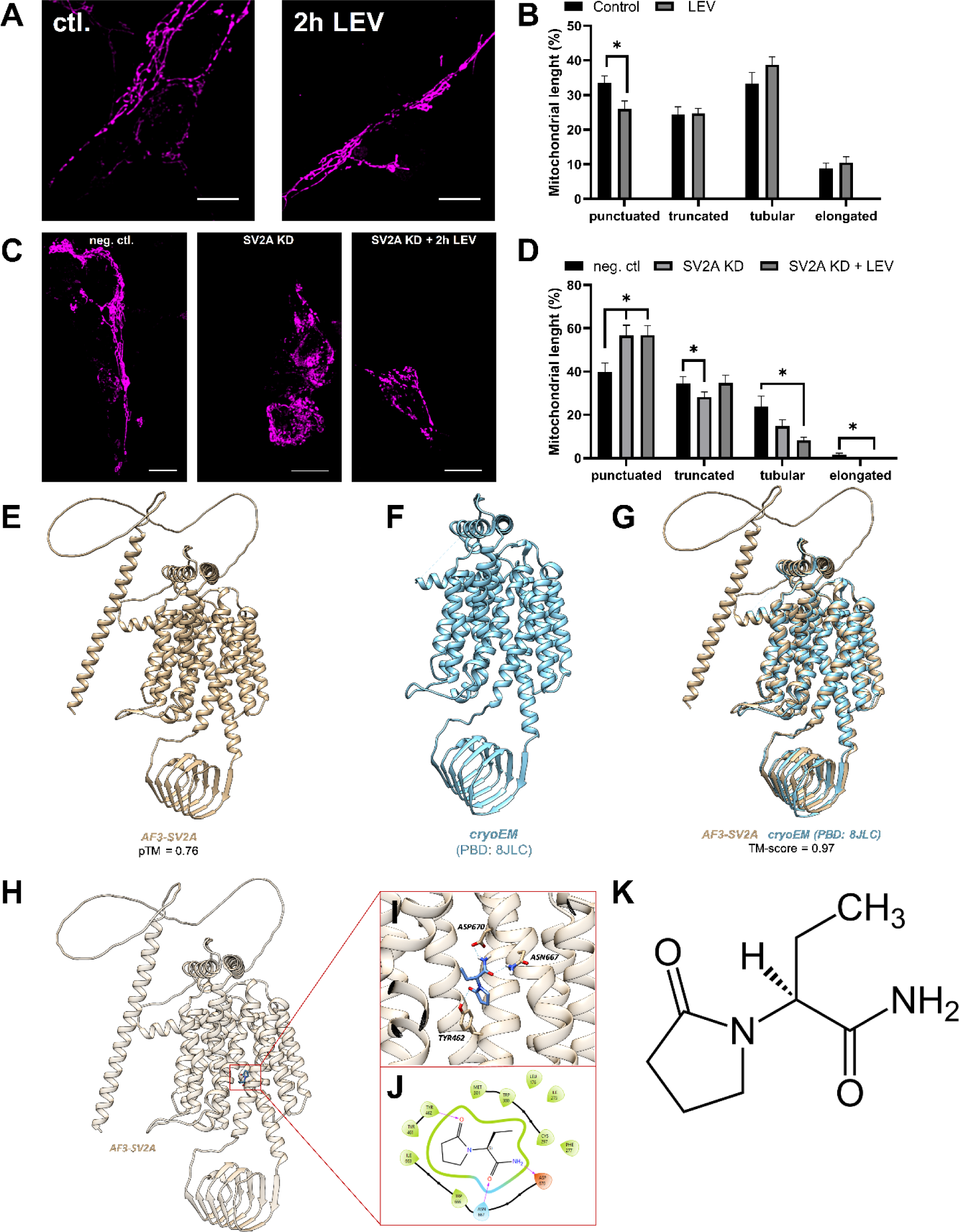
LEV treatment leads to elongated mitochondria in SH-SY5Y cells. A Representative confocal fluorescence images of mitochondrial morphology in levetiracetam-treated human SH-SY5Y cells (200 µM/2 h). Scale bar: 10 µM. B Length distribution of mitochondria in levetiracetam-treated cells compared to untreated control. Levetiracetam treatment elongates mitochondria in SH-SY5Y cells. C Representative confocal fluorescence images of mitochondrial morphology in human SH-SY5Y cells treated with scrambled siRNA (10 nM/48 h), SV2A siRNA (10 nM/48 h), and SV2A siRNA (10 nM/48 h) along levetiracetam-treatment (200 µM/2 h). Scale bar: 10 µM. D Length distribution of mitochondria in siRNA-induced SV2A KD cells treated with levetiracetam (200 µM/2 h) compared to siRNA-induced SV2A KD-only cells and negative control. The effect of levetiracetam on mitochondria vanishes during siRNA-induced SV2A KD. E Predicted structure of AF3-SV2A with corresponding pTM-score. F cryo-EM structure of SV2A (PDB: 8JLC). G Superposition of AF3-SV2A (gold) and the cryo-EM structure (blue) (PBB:8JLC) with corresponding TM-score. H-J Induced Fit calculation of AF3-SV2A with LEV. H Structure of AF3-SV2A with LEV after the IDF calculation. I Binding situation of LEV within AF3-SV2A. J Interaction diagram of LEV within AF3-SV2A. K Chemical structure of LEV. Data information: In (A-D) n=9-11 independent biological experiments. Data is expressed as mean ± SEM; student’s unpaired t-test (*p <0.05).

In this study, we employed the latest AlphaFold3 server to predict the three-dimensional structure of the synaptic vesicle protein 2A (SV2A), referred to as AF3-SV2A (Fig.3E-G). The accuracy of the predicted structure was evaluated using the predicted template modeling (pTM) score, an extension of the template modeling (TM) score. The TM score is a well-established metric that assesses the accuracy of predicted protein structures^42,43^. A pTM score greater than 0.5 suggests that the predicted fold is likely to be similar to the actual structure. The AF3-SV2A model exhibited a pTM score of 0.76, indicating a highly reliable predicted structure. Subsequently, the AF3-SV2A model was aligned with the recently resolved cryo EM structure of SV2A (PDB ID: 8JLC)^44^. The alignment yielded a TM score of 0.97, signifying an almost identical structural conformation between the predicted and experimental models. This high TM score underscores the exceptional predictive accuracy of AlphaFold3 for the SV2A protein.

To further validate the predicted model, we conducted induced fit docking calculations using the AF3-SV2A structure and LEV (Fig. 3H). The docking results showed a docking score of -8.420. Compared to the cryo-EM structure of LEV bound to SV2A it was observed that both identical binding partners. Specifically, LEV formed hydrogen bonds with residues THR462, ASP670 and ASN667 in both the predicted and experimental complexes, demonstrating the predictive accuracy of the AF3-SV2A model in replicating the binding interactions seen in the cryo-EM data^44^. Our findings indicate that the AlphaFold3-predicted structure of SV2A not only closely mirrors the experimentally determined conformation but also accurately predicts its binding interactions, thereby providing a robust model for further studies on SV2A’s functional interactions and mechanisms.

### Autophagy is reduced in SV2A KD SH-SY5Y cells

Since SV2A knockdown impairs mitochondrial morphology, we further investigated whether SV2A affects mitochondrial length distribution by reducing punctuated mitochondria via autophagy. First, we examined whether SV2A itself has a direct impact on autophagy by performing siRNA-mediated SV2A knockdown and additionally treating the cells with 600 nM V-ATPase inhibitor bafilomycin A1 to block the lysosome-mediated degradation of autophagosomes. Autophagic flux was determined by Western blotting, subtracting the LC3BII band intensity of samples with intact autophagy from those treated with bafilomycin A1, with LC3BII serving as a marker for autophagy. We observed a significant reduction in autophagic flux in SV2A knockdown cells (Fig. 4A).

**Figure 4.**
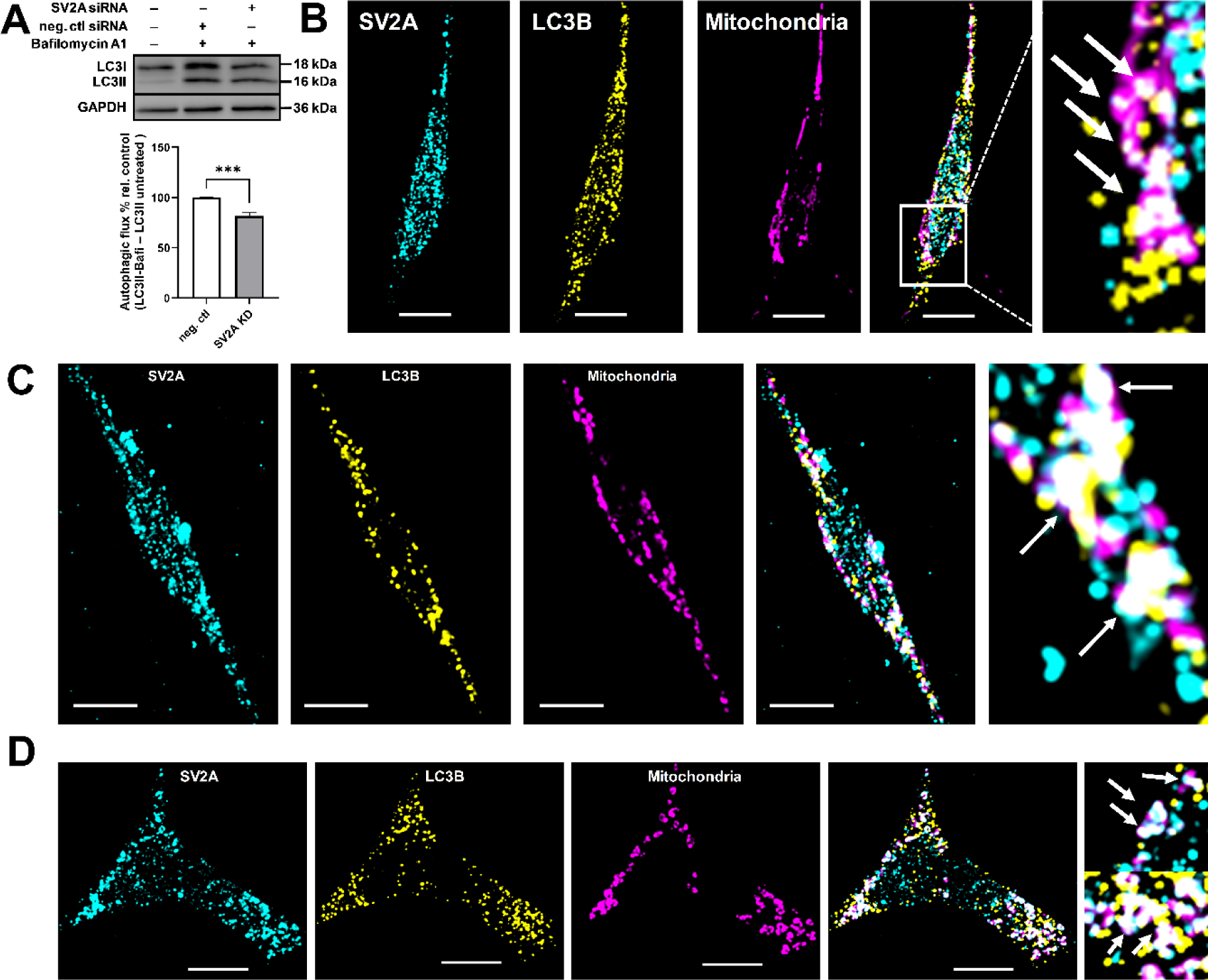
The role of SV2A in autophagy and mitophagy. A siRNA mediated SV2A knockdown leads to reduced autophagic flux. Representative LC3B western blot of SV2A KD and neg. ctl SH-SY5Y cells treated with bafilomycin A1 (600 nM/4 h). GAPDH serves as an internal control. 1. well: control cells neither treated with bafilomycin A1 nor with siRNA, 2. well: cells treated with scrambled siRNA (10 nM/48 h) and bafilomycin A1 (600 nM/4 h), 3. well: cells treated with SV2A siRNA (10 nM/48 h) and bafilomycin A1 (600 nM/4 h). Autophagic flux in SV2A KD cells compared to cells treated with scrambled siRNA. B Colocalization of SV2A with the autophagy marker LC3B. Representative confocal images of SH-SY5Y cells stained for SV2A, LC3B and mitochondria treated with bafilomycin A1 (600 nM/4 h). Colocalization of the three is indicated by bright white spots. Scale bar: 10 µm. C Representative confocal images of CCCP (10 µM/1 h) and bafilomycin A1 (600 nM/4 h) treated SH-SY5Y cells stained for SV2A, LC3B and mitochondria. Colocalization of the three is indicated by bright white spots. Scale bar: 10 µm. D Representative confocal images of rapamycin (10 µM/4 h) and bafilomycin A1 (600 nM/4 h) treated SH-SY5Y cells stained for SV2A, LC3B and mitochondria. Colocalization of the three is indicated by bright white spots. B-D Enlarged images of boxed areas are displayed at the end of each image series. Scale bar: 10 µm. Data information: In (A) n=5 independent cell extracts. In (B) n=18 independent biological experiments. In (B-D) Control: n=18; CCCP treated: n=11, rapamycin treated: n=13. Data are expressed as mean ± SEM; student’s unpaired t-test (*p <0.001).

Second, we investigated how SV2A might modulate autophagy. We utilized confocal microscopy to analyze the potential colocalization of SV2A and LC3B in SH-SY5Y cells treated with 600 nM bafilomycin A1. Therefore, we determined SV2A, LC3B, and mitochondrial localization (Fig. 4B) with individual analyses performed for SV2A and mitochondria, SV2A and LC3B, and LC3B and mitochondria. We observed colocalization of SV2A with mitochondria, as expected confirmed by a Pearson’s correlation coefficient of 0.74 (Figure 5C). Furthermore, SV2A and LC3B were found to colocalize at mitochondria and in the cytosol, although LC3B was often observed apart from SV2A (Figure 5D). The colocalization of SV2A and LC3B was further supported by a Pearson’s correlation coefficient of 0.76 (Figure 5D). Additionally, consistent with its role as an autophagosomal protein, LC3B was frequently detected at mitochondria, indicative of its involvement in mitophagy (Figure 5E). The Pearson’s correlation coefficient of 0.74 further supports the localization of LC3B at mitochondria (Figure 5E).

**Figure 5.**
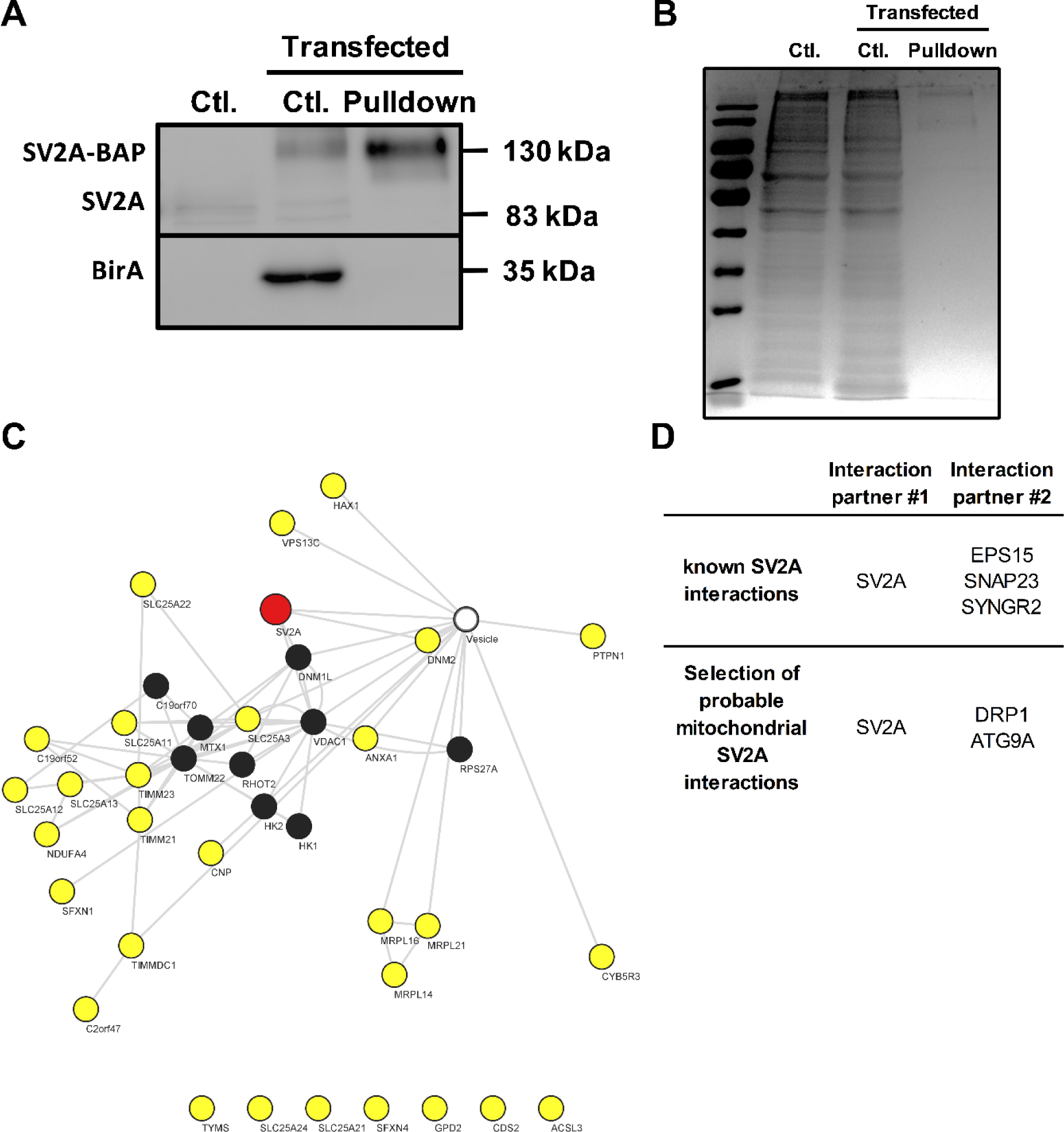
SV2A pulldown reveals interaction with mitochondrial and vesicle. A Purification of SV2A in HEK293 cells coexpressing SVA2 and BirA. As shown SV2A can be effectively purified and enriched in the eluate described in Material and Methods. SV2A-BirA construct band is seen at around 130 kDa. Native SV2A band was detected at 83 kDa. The lower panel shows that no purified SV2A is detected in the absence of BirA. n=3. B Preparative purification of SV2A in a 12% SDS-PAGE. C SV2A-interacting proteins in HEK293 cells identified by reversible cross-link immuno-precipitation (ReCLIP) LC-MS analysis. Network analysis was conducted using String (http://string-db.org/, version 12.0, String default settings). The network was visualized using the Cytoscape program and the strong default setting for confidence assignment. SV2A is highlighted in red color. Proteins of the outer mitochondrial membrane are indicated in black and proteins of the mitochondria in general (including inner mitochondrial membrane proteins) in yellow colors. The white circle summarizes all vesicle proteins. D Identification of known and probable interactions of SV2A predicated on String program determination and LC/MS analysis. Data information: In (A-D) n=3 independent biological experiments.

Third, to determine whether alterations in SV2A and LC3B are directly linked to the induction of autophagy or more specifically mitophagy, we treated our SH-SY5Y cells with 10 µM of the mitochondrial uncoupler CCCP (inducing mitophagy) or 10 µM rapamycin (inducing autophagy), along with additional bafilomycin A1 treatment to inhibit autophagosomal degradation. For mitophagy, we observed an increase in the colocalization of SV2A, LC3II, and mitochondria under mitochondrial stress compared to control cells treated with only of SV2A, LC3B, and mitochondria during mitophagy induction by CCCP, compared to control cells treated with bafilomycin A1 only (Fig. 4C). This may indicate that SV2A increasingly localizes to dysfunctional mitochondria during mitophagy induction. Similar to control cells, SV2A and LC3B were found to colocalize at mitochondria and in the cytosol, but to a greater extent at mitochondria (Fig. 4C).

After rapamycin-induced autophagy, we observed an altered distribution of SV2A. During autophagy induction, there was an increase in the colocalization of SV2A, LC3B, and mitochondria (Fig. 4D). SV2A becomes predominantly concentrated at the cell periphery, where fragmented mitochondria accumulate, forming large agglomerates on the mitochondria themselves (Fig. 4D). Additionally, similar to mitophagy, we observed colocalization of SV2A and LC3B at both mitochondria and in the cytosol, with a higher abundance at mitochondria (Fig. 4D). Autophagy induction leads to the localization of LC3B primarily to severely fragmented mitochondria, resembling CCCP-treated cells (Fig. 4D).

Our results suggest that SV2A might play a pivotal role in autophagy regulation, evidenced by its influence on mitochondrial morphology, localization to dysfunctional mitochondria during stress-induced mitophagy, and alterations in autophagic flux.

### Identification of mitochondrial proteins as interaction partners of SV2A

Since other vesicular proteins such as SYN17 mediate their mitochondrial effects via the interaction with mitochondrial proteins, we decided to test our third hypothesis identifying protein-protein interactions using reversibly cross-linked immunoprecipitation (ReCLIP) followed by mass spectrometry. To biotinylate SV2A, a 15-amino acid biotin acceptor peptide (BirA) sequence was added to the C-terminal region of SV2A full length. The resulting construct was coexpressed in HEK293T cells along with BirA, a bacterial protein-biotin ligase. HEK293T cells were chosen due to their low endogenous SV2A protein levels. Subsequently, we determined whether SV2A was efficiently biotinylated. As anticipated, the antibody detected a band at 37 kDa, corresponding to the endogenously expressed SV2A protein in non-transfected control cells (Fig. 5A). When cells were co-expressed with the biotinylated SV2A (SV2A-BAP) construct and BirA, another band, which shifted to a higher molecular weight (130 kDa), was observed via Western blot (Figure 8A). In contrast, no shifted band was observed when only SV2A-BAP or BirA were expressed in transfected control cells (Fig. 5A). The efficiency of biotinylation was verified by binding tagged SV2A in crude cell extracts to streptavidin-coupled paramagnetic beads. Western blot analysis of the material eluted from the beads showed that tagged SV2A protein was enriched in the transfected pulldown samples (Fig. 5A). The purified extracts were separated using SDS-PAGE (Fig. 5B).

After isolating the pulldown samples treated either with or without RNase, along with control pulldowns from singly transfected cells (either SV2A-BAP or BirA), eluted proteins were digested with trypsin. The tryptic peptides were separated by nanoUPLC directly coupled to a Synapt G2-S mass spectrometer operated in ion-mobility-enhanced data-independent acquisition mode. Overall, we identified and quantified over 220 proteins with less than 1% false discovery rate (FDR) (Supplementary Table 1, Supporting Information). 143 proteins were specifically associated with SV2A, as determined by our mass spectrometry analysis. Among these, 113 proteins were detectable only in pulldowns from double-transfected cells, while an additional 30 proteins were found to be at least 2-fold more abundant compared to controls. The high proportion of proteins detected exclusively in the pulldown samples confirms the high specificity of the ReCLIP approach. We identified several mitochondrial protein interaction partners, as well as a plethora of vesicular proteins interacting with SV2A (Fig. 5C,D). The most interesting interaction partners were the mitochondrial fission factor DRP1 and the autophagy protein ATG9A regarding SV2A effects on mitochondrial morphology and autophagy.

### SV2A and DRP1 interact at mitochondria

Having detected a protein-protein interaction between SV2A and DRP1, our next objective was to visualize this interaction at the mitochondrial level. Since DRP1 translocates from the cytosol to the outer mitochondrial membrane (OMM) to induce mitochondrial fission, we investigated the interaction between SV2A and DRP1 after inducing mitochondrial fission using the complex I inhibitor rotenone, employing both confocal microscopy and super-resolution microscopy (Fig. 6). SH-SY5Y cells were treated with 5 µM rotenone for 24 h. As expected, rotenone induced mitochondrial fragmentation and the translocation of DRP1 to mitochondria (Fig. 6A). Additionally, we observed an increase in SV2A colocalization with fragmented mitochondria. The increased colocalization of SV2A and mitochondria caused by mitochondrial stress was confirmed by an increase in Pearson’s correlation coefficient from 0.78 to 0.84 compared to control cells. Colocalization of SV2A and DRP1 increased during rotenone treatment and was almost exclusively restricted to mitochondria (Fig. 6A). Accordingly, Pearson’s correlation coefficient (r) for SV2A and DRP1 rose from 0.78 to 0.82 compared to the control.

**Figure 6.**
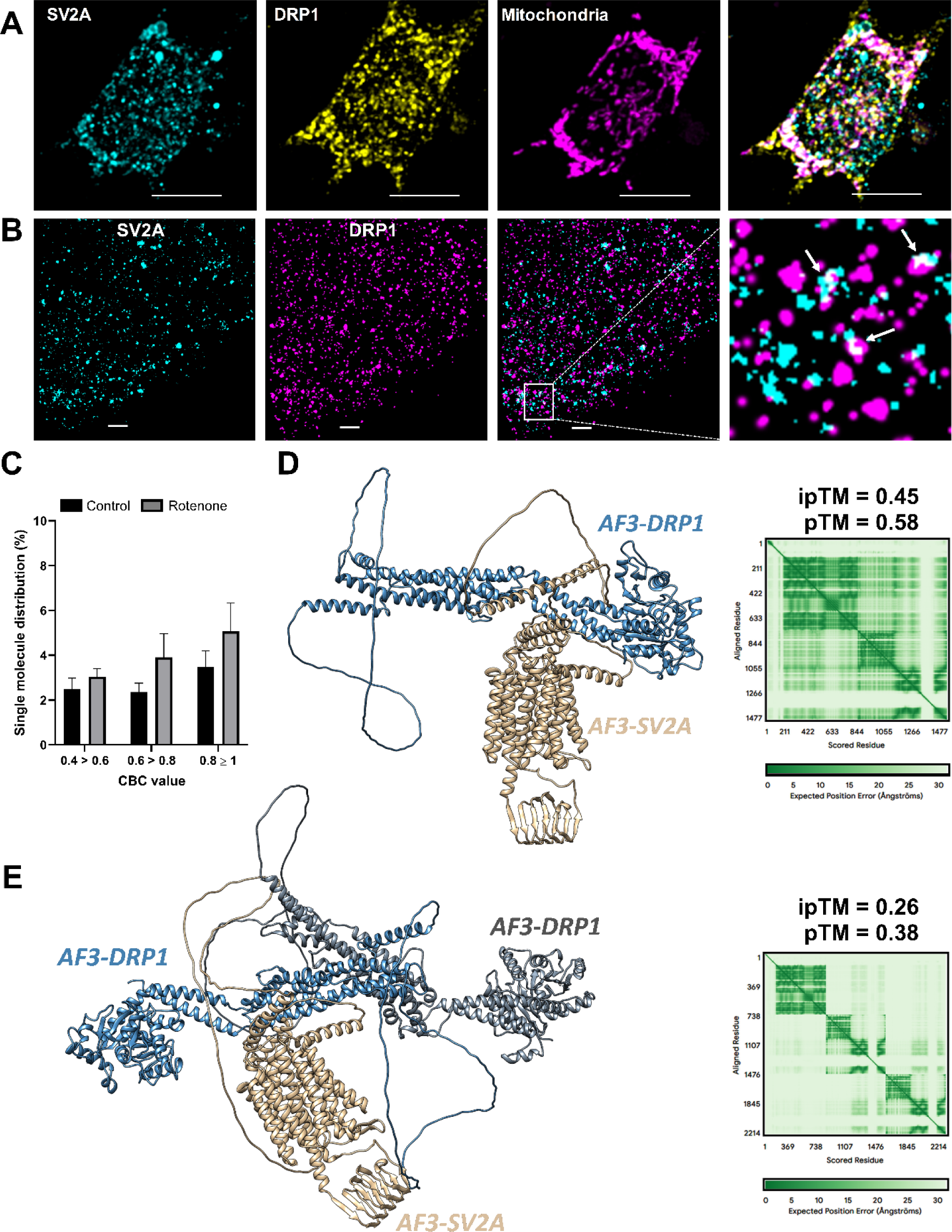
SV2A and DRP1 interaction. A Representative confocal fluorescence images of rotenone-treated (5 µM/24 h) human SH-SY5Y cells stained for SV2A, DRP1, and mitochondria. Colocalization of the three proteins of interest is indicated by bright white spots. Scale bar: 10 µm. B Representative super-resolution images of human SH-SY5Y cells treated with rotenone (5 µM/24 h) stained for SV2A and DRP1. Scale bar: 1 µm. C Histogram of CBC value distribution for the colocalization of SV2A and DRP1 in control and rotenone-treated cells. Enlarged are CBC values corresponding to the colocalization of the two proteins. D Predicted model of SV2A interacting with a DRP1 dimer by AlphaFold3 with corresponding ipTM and pTM values. E Predicted model of SV2A interacting with a DRP1 monomer by AlphaFold3 with corresponding ipTM and pTM values. Data information: In (A) Control: n=16 independent biological experiments; n=11 rotenone treated. In (B) Control: n=9; rotenone treated: n=9. Data are expressed as mean ± SEM; student’s unpaired t-test (*p <0.05; **p <0.01, ***p <0.001).

In summary, the confocal data confirmed that mitochondrial stress increases the mitochondrial localization of both SV2A and DRP1, resulting in increased colocalization of these two proteins. To validate this observation, we further analyzed rotenone-treated SH-SY5Y cells using super-resolution microscopy (Fig.6B,C). The cells were immunostained for SV2A with ATTO488 and for DRP1 with Alexa647. The super-resolution data confirmed increased colocalization of SV2A and DRP1 under rotenone treatment. When SV2A molecules with a CBC ≥ 0.5 were displayed, 12.8% of SV2A and DRP1 molecules colocalized with each other (Fig. 6C).

Since the AlphaFold3 predicted model of SV2A showed high accuracy and reliability, we used AlphaFold3 to predict the structural conformation and interactions between SV2A and DRP1. We started with the interaction between SV2A and a DRP1 monomer (Fig 6D). The prediction resulted in an interface predicted TM-score (ipTM) of 0.45 and a predicted TM-score (pTM) of 0.58. Typically, a high-confidence prediction requires ipTM values greater than 0.5 and pTM values greater than 0.6. Although our predicted values are close to these thresholds, they fall slightly short, indicating moderate confidence in the predicted interaction.AlphaFold3’s prediction suggested that the cytoplasmic N-Terminus of SV2A (11 - 26, 95 – 106) binds to the BSE region of DRP1 (307-318) and the G Domain region. In addition, AlphaFold3 suggests an interaction between cytoplasmic SV2A domain (360 – 401; 406 – 420) with the BSE region (290 – 320) and the G Domain region (200 – 299) of DRP1 respectively. However, it is important to note that this prediction has limitations and should be interpreted with caution.

Since DRP1 mainly exists as dimers or higher oligomers^9–11^, we predicted the interaction of SV2A with two DRP1 molecules to simulate DRP1 dimerization (Fig 6E). In this scenario, the ipTM value was 0.26 and the pTM value was 0.38, indicating a lower confidence in the predicted interaction. Despite this, the dimerization of DRP1 was predicted correctly, with both DRP1 molecules interacting via the stalk region, which has been experimentally validated previously. Interpreting these values, the moderately confident ipTM and pTM scores from the SV2A-DRP1 prediction suggest a potential interaction site between the cytoplasmic SV2A domain (364-421) with the MD stalk domain of DRP1 (359 – 388 and 419 – 462). In addition, an additional interaction was proposed between the cytoplasmic N-Terminus of SV2A (91-97) with the stalk region of DRP1.

## Discussion

It is well-established that SV2A is a crucial protein for vesicle function, changes, and distribution in brain vesicles and synapses. Other functions outside of vesicles are not well described until today. Here we report three novel findings using a broad set of methods ranging from imaging to proteomics and AI driven protein modelling which suggest a completely new role of SV2A apart from vesicles. First, we decipher the mitochondrial localization of SV2A at the OMM and demonstrate that inactivation of SV2A either in a siRNA mediated SV2a KD in cells or a SV2a KO mouse model, leads to mitochondrial fragmentation. Secondly, SV2A also affects the basal autophagic flux as well as mitophagy. Thirdly, we identify several mitochondrial protein interaction partners as well as a plethora of vesicular proteins interacting with SV2A, using ReCLIP followed by mass spectrometry. The interaction of SV2A with the fission factor DRP1 and the autophagy protein ATG9A might explain the effects of SV2A on mitochondrial morphology and autophagy. Our findings with AlphaFold3 provide first insights into the potential molecular interactions between SV2A and DRP1. These results suggest that SV2A might interact at OMM with DRP1 and thereby inhibit DRP1 oligomerization and DRP1 interaction with mitochondrial fission receptors such as MID49 both necessary for mitochondrial fission.

Many studies have traditionally stated SV2A solely as a vesicular protein. However, at the mitochondrial level, our group was the first to reveal mitochondrial effects of the SV2A ligand LEV^45^. Here, we demonstrate for the first time the precise subcellular localization of SV2A at mitochondria, suggesting its specific compartmentalization within the mitochondrial structure at OMM using confocal microscopy and super-high-resolution microscopy GSIDM. Our data are supported by another study from Kislinger et al., which revealed that SV2A exhibits a 22-fold higher expression level in brain mitochondria compared to microsomes and the cytosol, using proteomic and transcriptomic analyses of brain organelles in 8-10-week-old female ICR mice^46^. Additionally, Graham et al. observed SV2A to be increased in synaptic mitochondria compared to non-synaptic mitochondria which further supports our hypothesis of SV2A being present at the mitochondrial level^47^. The authors suggest that this finding might be due to impurity issues of mitochondrial isolation techniques. With our current data, we assume that these results cannot be attributed to impurity issues of the mitochondrial isolation technique, but rather that SV2A is also localized at OMM.

Until now, the role of SV2A in influencing mitochondrial morphology and autophagy was not known. In the current study, we used a siRNA-mediated SV2A KD cell model and SV2A KO animals to show, that reduction in SV2A protein expression levels or a complete loss of SV2A is associated with alterations in mitochondrial morphology. Mitochondria are fragmented in SV2A KD cells and in CA3 neurons of SV2A KO animals compared to their respective controls. These results are supported by the effects of the SV2A ligand LEV, which leads to a significant increase in tubular mitochondria. These findings are further supported by other studies demonstrating that LEV specifically protects mitochondria and upregulates TOMM20 in a spinal muscular atrophy iPSCs model^48^ improves mitochondrial membrane potential in C17.2 neuronal cells^49^ and protects against mitochondrial dysfunction and oxidative stress in a status epilepticus rat model^50^.

There are two possible explanations for our findings so far. First, SV2A KD influences mitochondrial fission and fusion directly. Second, SV2A interacts with other proteins involved in mitochondrial dynamics such as DRP-1 which was already reported for other vesicular proteins. In the last years, there has been growing interest in the participation of SNARE proteins regulating mitochondrial fission and fusion as well as autophagy and in particular mitophagy^51,52^. The vesicle protein SYN17 is also a mitochondria-associated protein implicated in mitochondrial dynamics promoting mitochondrial fission in fed cells via regulating DRP1 localization and activity and autophagy in starved cells switching its binding from DRP1 to ATG14L inhibiting autophagy^35^. In contrast, Stx4 enrichment at mitochondria favors mitochondrial fusion^36^. Regarding the effects on autophagy, SYN17 and C. elegans SNARE proteins such vamp-7 share similar effects on autophagy ^53^. In contrast, we observed a significant decrease in autophagic flux upon SV2A KD in SH-SY5Y cells suggesting that SV2A improves autophagy/mitophagy. But not only SNARE proteins are involved in autophagy or mitophagy but also mitochondrial fission and fusion factors such as DRP1 or MFN-2^54–56^.

Therefore, we asked whether this might also be true for SV2A. For this experiment, we consciously used HEK293T cells because of the low endogenous SV2A expression within this model and analyzed them by mass spectrometry. Overall, we identified and quantified >220 proteins, whereas 143 are found to be specifically associated with SV2A itself. As expected we observed several vesicle proteins to interact with SV2A such as SNAP-23 and synaptogyrin-2. Importantly, we also found several mitochondrial proteins to interact with SV2A such as the fission factor DRP1 and ATG9A, the only transmembrane protein of the autophagy network. ATG9A is a lipid scramblase known to enable lipid transfer to the phagophore membrane during early stages of autophagy^14,57^. Our data regarding the role of SV2A suggests that SV2A might also be involved in the early stages of autophagy. The interaction between SV2A and ATG9A may indicate that SV2A acts as a binding partner for ATG9A containing vesicles and might be involved in phagophore expansion^14,57^. Knocking down SV2A (SV2A KD) may lead to increased mitochondrial fragmentation, as ATG9A loses one of its mitochondrial binding partners leading to reduced phagophore formation and autophagic flux.

Finally, we provide mechanistic data showing the potential molecular interactions between SV2A and DRP1 using Alphafold 3. It predicts interactions between the cytoplasmic N-Terminus of SVA2 and the BSE domain and the G domain of the DRP1 monomer as well as the cytoplasmic domain of SV2A with the BSE domain and the G domain. The cytoplasmic domain of SV2A is predicted to also interact with MD stalk domain of DRP1 which is considered to be important for the self-assembly into dimers, tetramers and higher order oligomers^11^. Especially the computed interaction of SV2A with conserved L1N^s^ loop at the receptor interface 3 of the MD DRP1 domain is of high interest as it is the site of the co-assembly with the mitochondrial fission receptor MID49^9,10^. In conclusion, while AlphaFold3 provides valuable predictions about protein interactions, these findings come with limitations. The moderate confidence levels necessitate further experimental validation to confirm the interaction dynamics and functional implications of SV2A and DRP1, particularly in the context of DRP1 dimerization and possible effects of SV2A shielding DRP1 from binding to mitochondrial fission receptors such as MID49 and thereby affecting mitochondrial morphology.

Together, these results prompt us to conclude that SV2A is not only a vesicular protein, as it is also located at the OMM regulating mitochondrial morphology as well as autophagy. SV2A seems to be an adaptor protein at the OMM which interacts with DRP1 inhibiting its oligomerization and interaction with mitochondrial fission receptors resulting in improved mitochondrial morphology. On the other hand, mitochondrial SV2A localization is increased under mitochondrial stress reflected in mitochondrial fragmentation. Under these conditions, SV2A co-localization with LC3B and DRP1 is also elevated. We propose that under mitochondrial stress, SV2A might induce autophagy by interacting with ATG9A.

## Methods

### Cell Culture

SH-SY5Y cells were cultivated in high-glucose Dulbecco’s modified Eagle’s medium (DMEM) (Gibco™, 10569010) supplemented with GlutaMAX™, 10% heat-inactivated fetal bovine serum (FCS) (Gibco^TM^, 10270-106), 60 units/mL penicillin, and 60µg/mL streptomycin (Gibco^TM^, 15140-122), 1% MEM vitamin solution (100x) (Gibco^TM^, 11120037), 1% MEM non-essential amino acids (100x) (Gibco^TM^, 11140035) and 1mM sodium pyruvate (Thermo Scientific™, 11360039) in a humidified incubator at 37°C containing 5% CO_2_.

Untransfected HEK293 cells were cultured in DMEM (Gibco^TM^, 41965-039) supplemented with 10% heat-inactivated FCS (Gibco^TM^, 10270-106) 50 Units/mL penicillin and 50 µg/mL streptomycin (Gibco^TM^, 15140-122), at 37 °C in a humidified incubator containing 5% CO_2_.

### Animals

Hippocampus-specific SV2A KO mice, in which SV2A is only knocked out in glutamatergic neurons of the dentate gyrus and CA3 region, were bread according to Menten-Dedoyart^41^. Animal care was in accordance with the declaration of Helsinki and the guidelines of the Belgium Ministry of Agriculture, in agreement with the European Community laboratory animal care and use regulations. Experimental research on the animals was performed with the approval of the University of Liège ethics committee (Belgium), filed under numbers 1258 and 1753, and accepted in 2011 and 2016, respectively.

### Confocal and super-resolution microscopy

5×10^5^ SH-SY5Y cells were seeded in full medium on glass coverslips for confocal or on µ-Dish 35 mm, high Glass Bottom dishes (Ibidi®, 81158) for super-resolution stainings, grown for 48 hours (37°C, 5% CO_2_) and later fixed with 4% paraformaldehyde (PFA) solution for 20 min at room temperature (RT). To quench leftovers of active PFA, samples were treated with quenching buffer (50 mM Tris HCl pH 8.0 and 100 mM NaCl in PBS) for 5 min followed by permeabilization with 0.2 %Triton X in phosphate buffered saline (PBS) for 20 min at RT. Afterwards, cells were washed with PBS for 5x5 min and blocked with 5% bovine serum albumin (BSA) in PBS for 1 hour at RT. Primary antibodies were incubated for 1 hour at RT. Samples were then washed with PBS and incubated with secondary antibodies for 1 hour at RT. For confocal imaging, mitochondria were stained with 200 nM MitoTracker^TM^ Deep Red FM (Invitrogen^TM^, M22426) for 45 min at 37°C before fixation. SV2A was stained with an anti-SV2A monoclonal antibody (Synaptic Systems; 119 011) followed by an secondary anti-mouse Alexa Fluor® 488 antibody (Abcam; ab150117) each for 1 hour at RT. For super-resolution microscopy, mitochondria were stained with anti-TOMM20-Alexa Fluor® 647 monoclonal Systems; 119 011) followed by a secondary anti-mouse ATTO488 antibody (NanoTag Biotechnologies; N1202-At488-S) each for 1 hour at RT.

After washing with PBS, coverslips for confocal microscopy were finally mounted onto slides with ProLong^TM^ Glass Antifade Mountant with NucBlue^TM^ Stain (Invitrogen™, P36985), and super-resolution samples were covered in PBS. Confocal images were acquired with the 63x oil immersion objective on the Leica TCS SP5 confocal microscope (Leica Microsystems GmbH, Germany) and super-resolution images with the 60 x 1,47 oil immersion objective of the Leica GSD 3D widefield super-resolution microscope (Leica Microsystems GmbH, Germany). Before imaging samples using the Leica GSD 3D widefield super-resolution microscope, fluorescent TetraSpeck^TM^ microspheres (Invitrogen^TM^, T7279) were measured for drift correction. A 1:500 TetraSpeck^TM^ dilution embedded in pyranose oxidase buffer + MEA was measured to determine the shift between green (488 nm) and red (647 nm) channels. This step is especially important for colocalization experiments as even a slight shift of a few nanometers between the channels alters the correct localization of the stained proteins which may lead to false results. After measuring TetraSpeck^TM^ beads, samples were embedded in pyranose oxidase buffer + MEA and ∼10,000 frames per channel were recorded. At the end of the session, TetraSpeck^TM^ beads were remeasured.

Fiji software was used to analyze images. To mimic oxidative stress, SH-SY5Y cells were treated with complex I inhibitor rotenone (Sigma-Aldrich_®_, R8875) 24 h before fixation and to assess the effects of LEV on mitochondria, cells were incubated with 200 µM levetiracetam 2 h before fixation.

### Microscope data processing and analyzation

Colocalization of proteins in confocal images was analyzed both visually on merged images and by Pearson’s correlation coefficient (Pearson’s r) calculated using the Fiji plugin JACoP. Pearson’s r measures the linear correlation between two datasets ranging from -1 (100% anti-correlation) to 1 (100% correlation). As SRM raw data consists of thousands of frames per channel containing the coordinates of stained proteins, ThunderSTORM (Fiji plugin) was used to extract the protein coordinates, process them (e.g. shift correction), and finally transform them into an image. Shift during the acquisition was calculated and adjusted for the red channel with custom-written MATLAB software (Dr. Márton Gelléri) using the data from the TetraSpeck^TM^ beads measurements. Transformed SRM images were then analyzed both visually on merged images and with Thunderstorm’s colocalization tool Coordinate-based colocalization (CBC). The CBC value is calculated for each coordinate over the distance between two coordinates in a 50 nm radius and ranges from -1 (100% anti-correlation) to 1 (100% correlation).

### Prediction of mitochondrial localization

To predict mitochondrial subcellular localization, we used the SV2A protein sequence in fasta format. The fasta file was uploaded and analyzed using the iMLP Technical University of Kaiserslautern, Kaiserslautern, Germany) (http://imlp.bio.uni-kl.de/) (accessed on 29 April 2024) tool^58^. The SV2A fasta file was also uüloaded in DeepLoc 2.0 (DTU Health Tech, Lyngby, Denmark) (https://services.healthtech.dtu.dk/service.php?DeepLoc-2.0) (accessed on 29 April 2024).

### SV2A Knockdown

7×10^5^ SH-SY5Y cells were grown for 24 hours until 60-80% confluency. The next day, cells were treated with pre-designed Silencer^®^ Select SV2A siRNA (Thermo Scientific^TM^, s19182), Lipofectamine™ RNAiMAX (Invitrogen™, 13778075) and Opti-MEM^®^ Media (Gibco™, 11058021) according to the manufacturer’s instructions. As positive control siRNA stained with Alexa Fluor 555 was used to identify transfected cells. Cells were incubated with SV2A or negative control siRNA/Lipofectamine^TM^ RNAiMAX reagent for 48 hours.

### Measuring mitochondrial length

Mitochondrial length was measured on confocal images using FIJI software. During data analysis, 100 mitochondria were measured per n and divided into four groups according to Stockburger et al.^45^.

### Preparation of protein extracts

Cells were seeded 72 h in advance and synchronized with ice-cold PBS. For analysis cells were washed twice with PBS and lysed on ice in radio-immunoprecipitation assay buffer (RIPA) containing 50 mM Tris-HCl (pH= 7.4) (Carl Roth, 5429.5), 150 mM sodium chloride (Carl Roth, P029.3), 1% Triton™ X-100 (Carl Roth, 3051.4), 0.5% sodium deoxycholate (Sigma-Aldrich, 30970), 0.1 % sodium dodecyl sulfate (SDS) (Carl Roth, CN30.2), 5 mM ethylenediaminetetraacetic acid (Carl Roth, CN06.2) and 1 mM phenylmethylsulfonylfluoride (PMSF) (Carl Roth, 6367.1). After incubation with RIPA buffer for 45 min samples were centrifuged (10,000 g, 4°C, 10 min). Total protein concentration was estimated using Bradford method (Bio-Rad Protein Assay Dye Reagent Concentrate, Bio-Rad Laboratories, #5000006).

### Western Blot (WB)

20 µg protein per lane was loaded on a 12% SDS polyacrylamide gel electrophoresis (PAGE). Samples for SV2A analysis were not boiled. After separation, samples were transferred onto a polyvinylidene fluoride membrane (Thermo Scientific™, 88520) and incubated with 5% BSA for blocking of nonspecific binding at RT. All membranes were washed 3x with Tris buffered saline with Tween® solution (TBST) and then incubated with primary antibodies overnight at 4°C. Primary antibodies were purchased from Abcam, Cambridge, United Kingdom (Anti-SV2A, ab32942; Anti-α-tubulin, ab7291; Anti-DRP1, ab184247; Anti-GAPDH, ab181602; Anti-TOMM20, ab186735), Thermo Scientific™ (Anti-Adenylate kinase 2, MA5-29016; Anti-TIMM23, MA5-27384), and Sigma-Aldrich (Anti-ß-Actin, A1978, Anti-Citratsynthetase, SAB2702186, Anti-LC3B, L7543). After washing the membrane 3x with 0.5% TBST, membranes were treated with horseradish peroxide conjugated secondary antibodies. For Adenylate kinase 2, GAPDH, LC3II and SV2A an anti-rabbit antibody was used (Sigma-Aldrich, A0545), for α-Tubulin, β-actin, Citratsynthetase, and TIMM23 an anti-mouse-antibody was used (Invitrogen^TM^, 31430). Last, all membranes were washed 3x with 0.5% TBST and analyzed with Amersham ECL™ Prime Western Blotting Reagent (Cytiva, RNP2236) using a Fusion Pulse TS Imager (Vilber Lourmat, France). All bands were analyzed according to their appropriate kDa.

### Mitochondrial isolation

2.5×10^6^ HEK-293 cells were seeded in full medium and grown until they reached ∼90% confluency after ∼96 hours. When 90% confluency was reached, medium was aspirated, and cells were washed with PBS, collected, and centrifuged (700 g, 5 min, 4°C). The resulting pellet was then resuspended in mitochondrial isolation buffer (MIB) containing 400 mM Saccharose, 1M Tris/MOPS, 200 mM EGTA/Tris and broken up with a Potter S (B. Braun Biotech International, Germany). The disruption was continued by drawing the cell suspension into a syringe with an 18-gauge needle and expelling it with a 27-gauge needle. Drawing and expelling were repeated 5x. Broken-up cells were transferred to fresh tubes and centrifuged in a swinging bucket rotor (600 g, 5 min, 4°C). The resulting supernatant was carefully removed and centrifuged another time in a fixed-angle rotor (10,000 g, 5 min, 4°C). After centrifugation, the mitochondrial pellet was resuspended in HS buffer and mitochondrial concentration was determined by using Pierce™ BCA Protein Assay Kits (Thermo Scientific™, 23225) according to the manufacturer’s protocol.

### Mitochondrial fractioning and TCA protein precipitation

Mitochondrial fractioning was performed by incubating isolated mitochondria with increasing concentrations of digitonin, causing mitochondrial membranes to dissolve, depending on the digitonin concentration. Per sample 800 µg mitochondria were dissolved in 150 µL HS buffer and 150 µL digitonin solution. A total of thirteen samples were prepared, with the first two samples receiving no digitonin and the remaining eleven samples receiving increasing digitonin concentrations starting at 0.005% until a concentration of 0.105% was reached. Immediately after the addition of digitonin, proteinase K was added to all samples occasional inverting. To stop mitochondrial digestion, 100 mM PMSF was pipetted into each sample, incubated for 5 min, and centrifuged (13,000 g, 10 min, 4°C). After centrifugation, the pellet was resuspended in HS buffer/1 mM PMSF.

After successful mitochondrial fractioning, proteins were precipitated by trichloroacetic acid (TCA). 100% TCA solution was added to the samples and incubated for 10 min. After centrifugation (14,000 rpm, 5 min, 4°C), the pellets were washed with ice-cold acetone. After another centrifugation (14,000 rpm, 5 min, 4°C), the pellet was washed again with acetone. This procedure was repeated three times and afterwards, the pellet was dried on ice for 30 min. The dried pellet was then dissolved in 4x loading buffer by boiling the samples for 10 min at 95°C. The finished samples were stored at -80°C and later analyzed by WB. Marker proteins for each mitochondrial fraction were detected to determine in which mitochondrial fraction the protein of interest was located.

### Live cell imaging

5×10^5^ SH-SY5Y cells were seeded and grown for 24 hours. On the day of transfection, cells were treated with mEos2-Mito-7 plasmid, Silencer^®^ Select siRNA (Thermo Scientific^TM^, s19182) and Lipofectamine^TM^ RNAiMAX according to the manufacturer’s instructions. Cells were transfected with plasmid or siRNA/lipofectamine^TM^ RNAiMAX complex for 48 hours.

After successful transfection, live cell imaging was performed using the 100x oil immersion objective on the Visitron Spinning Disc microscope (Visitron Systems GmbH, Germany). During imaging, cells were kept at 37°C and 5% CO_2_ in a humified chamber and one image was acquired every 30 sec for a total duration of 10 min. Fiji software was used to analyze the recordings for fission/fusion events, distance traveled, and velocity. Fission and fusion events were determined visually for 10 minutes by comparing changes of mitochondria in each time frame to the next time frame.

### Histology of brain slices

#### Paraffin embedding

PFA fixed tissue was dehydrated by ascending ethanol dilutions, followed by xylene substitutes, and finally infiltrated with paraffin in the semiautomatic Leica Tissue Processor (Leica Microsystems GmbH, Germany). After 18 hours, the dehydrated and paraffin-infiltrated tissue was embedded in paraffin blocks using the Leica Tissue Embedding Center (Leica Microsystems GmbH, Germany). The finished paraffin blocks were stored at 4°C.

### Tissue sectioning

Paraffin blocks were trimmed and placed into the cooling specimen clamp of the Microtome (Leica Microsystems GmbH, Germany). Before cutting 10 µm thick slices, paraffin blocks were trimmed until the desired area was reached. During sectioning, the paraffin ribbon was placed into a water bath at 40°C to remove wrinkles, and sections were then mounted onto charged slides. Afterward, sections were dried for 30 min at 35°C on a heating plate and baked overnight at 47°C in an oven. Finished slides were stored at 4°C.

### Staining of brain slices

Brain sections were deparaffinized and rehydrated by incubation with xylene substitutes followed by descending ethanol dilutions. Afterward, antigen retrieval was performed by boiling the sections for 15 min at 95°C in TRIS-EDTA buffer. After cooling, the tissue was encircled by a hydrophobic pen to build a hydrophobic barrier necessary for applying reagents such as antibodies. Sections were washed with 0.1% Tween^®^/PBS buffer followed by PBS both for 5 min at RT. Permeabilization was performed by treating the samples with 0.5% Triton-X 100/PBS at RT for 1 hour. Samples were then washed with PBS and blocked with 3% BSA/PBS Buffer for 1 hour at RT. After blocking, the slices were incubated with primary antibody in a humified chamber overnight at 4°C. The next day sections were washed with PBS followed by a 1-hour incubation with a secondary antibody at RT. Finally, slides were washed with PBS and coverslips were mounted onto the sections using ProLong^TM^ Glass Antifade Mountant with NucBlue^TM^ Stain (Thermo Scientific^TM^, P36985). Slides were stored at 4°C and imaged using the 63x oil immersion objective on the Leica TCS SP5 confocal microscope (Leica Microsystems GmbH, Germany).

### Measuring of mitochondrial length in brain slices

Mitochondrial length was measured using Fiji software. From each mouse brain, z-stacks were acquired at three different locations in the CA3 region. Mitochondria were measured in three images per stack, as measurement of mitochondria in z-projects was not possible due to the large number of mitochondria.

### Reversible Cross-Link Immuno-Precipitation (ReCLIP) in HEK293 cells

For ReCLIP, 7×10^5^ HEK-293 cells were seeded. After 24h, HEK293 cells were transfected with 1 μg pcDNA3-cytBirA and 1 μg pcDNA3.1-SV2A-BirA-biotinylation-sequence plasmid for 48 hours. To allow in vivo biotinylation, biotin was added in excess to a final concentration of 150 μM. After successful transfection, protein interactions are stabilized using the cell-permeable, lysin-reactive crosslinker Dithiobis[succinimidiylproprionate] (DSP), allowing SV2A to be coimmunoprecipitated with its interaction partners. Cells were harvested in PBS, transferred to 1.5 mL tubes and crosslinking is performed by treating the cells with 0.75 mM DSP for 30 min at 37°C. Crosslinking is stopped by adding 1M Tris/HCl pH 7.4 and incubating the cells for 15 min at RT. To investigate if crosslinking was successful, we conducted WB. Therefore, samples are centrifuged (1000 U, 5 min, RT), supernatants are removed, and resulting pellets are resuspended in RIPA buffer. Protein extraction was performed as described previously. As the amide bonds of DSP-crosslinked proteins are thiol sensitive they can be cleaved by DTT and β-mercaptoethanol to serve as a control for successful crosslinking. To cleave DSP-related bonds, 1M DTT solution is applied to the DSP-treated protein extracts for 30 min at 37°C before loading the gel. For WB of DSP-treated samples, it is crucial to use a non-reducing loading buffer, otherwise β-mercaptoethanol will cleave the crosslinker bonds.

### Proteolytic digestion for mass spectrometric analysis

Samples were processed by single-pot solid-phase-enhanced sample preparation (SP3) as described before^59,60^. In brief, proteins bound to the streptavidin beads were released incubating the samples for 5 min at 95° in an SDS-containing buffer (1 % (w/v) SDS, 10 mM biotin in 10 mM Tris, pH 7.5). After elution, proteins and DSP cross-linker were reduced with 50 mM DTT for 30 min at 37 °C. Afterwards, temperature was increased to 45 °C and samples were incubated for another 10 minutes. Proteins were then alkylated for 30 min at room temperature using iodoacetamide (IAA). Excess IAA was quenched by the addition of DTT. Afterwards, 2 µl of carboxylate-modified paramagnetic beads (Sera-Mag Speed Beads, GE Healthcare, 0.5 μg solids/μl in water as described by Hughes et. al^59^ were added to the samples. After adding acetonitrile to a final concentration of 70% (v/v), samples were allowed to settle at RT for 20 min. Subsequently, beads were washed twice with 70 % (v/v) ethanol in water and once with acetonitrile. Beads were resuspended in 50 mM NH_4_HCO_3_ supplemented with trypsin (Mass Spectrometry Grade, Promega) at an enzyme-to-protein ratio of 1:25 (w/w) and incubated overnight at 37°C. After overnight digestion, acetonitrile was added to the samples to reach a final concentration of 95% (v/v) followed incubation at RT for 20 min. To increase the yield, supernatants derived from this initial peptide-binding step were additionally subjected to the SP3 peptide purification procedure^60^. Each sample was washed with acetonitrile. To recover bound peptides, paramagnetic beads from the original sample and corresponding supernatants were pooled in 2 % (v/v) dimethyl sulfoxide (DMSO) in water and sonicated for 1 min. After 2 min of centrifugation at 12,500 rpm and 4 °C, supernatants containing tryptic peptides were transferred into a glass vial for MS analysis and acidified with 0.1 % (v/v) formic acid.

### Liquid chromatography-mass spectrometry (LC-MS) analysis

Tryptic peptides were separated using an Ultimate 3000 RSLCnano LC system (Thermo Fisher Scientific) equipped with a PEPMAP100 C18 5 µm 0.3 x 5 mm trap (Thermo Fisher Scientific) and an HSS-T3 C18 1.8 μm, 75 μm x 250 mm analytical reversed-phase column (Waters Corporation). Mobile phase A was water containing 0.1 % (v/v) formic acid and 3 % (v/v) DMSO. Peptides were separated running a gradient of 2–35% mobile phase B (0.1% (v/v) formic acid, 3 % (v/v) DMSO in ACN) over 40 min at a flow rate of 300 nL/min. Total analysis time was 60 min including wash and column re-equilibration steps. Column temperature was set to 55°C. Mass spectrometric analysis of eluting peptides was conducted on an Orbitrap Exploris 480 (Thermo Fisher Scientific) instrument platform. Spray voltage was set to 1.8 kV, the funnel RF level to 40, and heated capillary temperature was at 275°C. Data were acquired in data-dependent acquisition (DDA) mode targeting the 10 most abundant peptides for fragmentation (Top10). Full MS resolution was set to 120,000 at *m/z* 200 and full MS automated gain control (AGC) target to 300% with a maximum injection time of 50 ms. Mass range was set to *m/z* 350–1,500. For MS2 scans, collection of isolated peptide precursors was limited by an ion target of 1 × 10^5^ (AGC target value of 100%) and maximum injection times of 25 ms. Fragment ion spectra were acquired at a resolution of 15,000 at *m/z* 200. Intensity threshold was kept at 1E4. Isolation window width of the quadrupole was set to 1.6 *m/z* and normalized collision energy was fixed at 30%. All data were acquired in profile mode using positive polarity. Samples were analyzed in three technical replicates.

### Data analysis and label-free quantification

DDA raw data acquired with the Exploris 480 were processed with MaxQuant (version 2.0.3)^61,62^, using the standard settings and label-free quantification (LFQ) enabled for each parameter group, i.e. control and affinity-purified samples (LFQ min ratio count 2, stabilize large LFQ ratios disabled, match-between-runs). Data were searched against the forward and reverse sequences of the human reference proteome (UniProtKB/Swiss-Prot, 20,361 entries, release April 2022) and a list of common contaminants. For peptide identification, trypsin was set as protease allowing two missed cleavages. Carbamidomethylation was set as fixed and oxidation of methionine as well as acetylation of protein *N*-termini as variable modifications. Only peptides with a minimum length of 7 amino acids were considered. Peptide and protein false discovery rates (FDR) were set to 1 %. In addition, proteins had to be identified by at least two peptides. Statistical analysis of the data was conducted using Student’s t-test, which was corrected by the Benjamini–Hochberg (BH) method for multiple hypothesis testing (FDR of 0.01). SV2A interacting proteins had to show a 2-fold enrichment as compared to the controls across all pulldown experiments.

### Modeling protein-protein interactions

#### SV2A-modeling

The webserver AlphaFold3 (https://golgi.sandbox.google.com/) was utilized to predict the protein-structure of SV2A. The amino acid sequence of SV2A was obtained from UniProt (Q7L0J3) and inserted into the webserver, with the molecule type defined as "protein." The resulting predictions were downloaded and analyzed using Chimera^63^.

### Protein structure preparation

Published crystal structures of DRP1 (PDB 4BEJ) and SV2A (PDB ID: 8JLC) were loaded into Maestro (Schrödinger Release 2022-4: Maestro, Schrödinger, USA, 2022). To prepare the protein for docking and simulations, the protein preparation wizard was used to assign bond orders, add hydrogens, create zero-order bonds to metals, create disulfide bonds, and fill in missing side chains and loops using Prime. Default parameters were used for the optimization of hydrogen-bond assignment.

### Induced Fit Calculation

Induced fit docking and ligand modeling of the interaction of LEV and AlphaFold3-SV2A was performed using Maestro (Schrödinger Release 2022-4: Maestro, Schrödinger, USA, 2021). AlphaFold3-SV2A was prepared before docking with the protein preparation wizard. In Maestro, original hydrogens were removed and replaced, bond orders were assigned, and the structure was minimized. A grid was prepared around the active site centered at the position of LEV. All ligands were prepared for docking in Maestro using the ligand preparation function. Ligands were docked to the active-site grid using Induced Fit Glide Docking with post-docking minimization. Visualization was processed with ChimeraX^63^.

### Setup and Building of the Systems

All MD simulations in this study were completed with the Desmond Molecular Dynamic software under Schrödinger (Schrödinger Release 2022-3: Desmond Molecular Dynamics System, D. E. Shaw Research, USA, 2022. Maestro–Desmond Interoperability Tools, Schrödinger, USA, 2022). Setups for the runs were assembled with Desmond System Builder application under Maestro. All simulations were run within an orthorhombic box full of explicit water molecules generated by the single-point charge (SPC) model^64^. The assembling was continued with water boxes on both the top and bottom of the membrane, also according to the buffer. The assembly was then completed with sodium and chloride ions to statistically reach the isotonic (0.15 M) concentration, and additional counter ions were added if needed to neutralize the charge of the peptide or conjugate so the net charge of the system was reduced to zero. Before the MD simulations, the assemblies were minimized with OPLS5 force field method for the final positioning of the molecules to avoid steric clashes.

### Molecular Dynamics

Desmond package in Schrödinger suite v2021-3 (Schrödinger Release 2022-3: Desmond Molecular Dynamics System, D. E. Shaw Research, USA, 2022. Maestro–Desmond Interoperability Tools, Schrödinger, USA, 2022) was used to run the MD simulation to elucidate the effectiveness of the screened compounds by molecular docking^65^. The ‘system builder’ was used to prepare the protein-ligand complex. The SPC water model in an orthorhombic shape was selected after minimizing the volume, with 10 Å × 10 Å × 10 Å periodic boundary conditions in the P-L complex’s x, y, and z-axis.

Further, using the OPLS2005 forcefield, the complex minimized its energies by heating and equilibrium processes before the production run of MD simulations^66^. Further, with the time step of 100 ps, the system normalized in an equilibrium state at 1000 steps. The final production run was kept for 20 ns, at the time steps of 100 ps, 300 K temperature and 1.01325 atm pressure, for both complexes applying the Nose-Hoover method with NPT ensemble^67^.

### Protein-Protein-Docking

The webserver AlphaFold3 (https://golgi.sandbox.google.com/) was utilized to predict the protein-protein interactions between SV2A and DRP1. The amino acid sequence of each protein was obtained from UniProt and inserted into the webserver, with the molecule type defined as "protein." The resulting predictions were downloaded and analyzed using Maestro and Chimera.

### Statistical analysis

Data are presented as mean ± S.E.M. For statistical comparison, student’s unpaired t-test was calculated with GraphPad Prism. *p* values of *p<0.05; **P<0.01, ***P<0.001 were considered statistically significant.

## Data availability

The mass spectrometry proteomics data have been deposited to the ProteomeXchange Consortium (http://proteomecentral.proteomexchange.org) via the jPOST partner repository ^68^ with the dataset identifiers PXD051373 (ProteomeXchange) and JPST003029 (jPOST). To review the data: Go to reviewer link https://repository.jpostdb.org/preview/126814650266180d867495c

Access key: 2024

## Acknowledgements

The contribution of CB and AK was supported by a grant from the Peter-Beate-Heller Foundation of the Deutsche Stifterverband.

## Author contributions

M.J., J.S.R., K.P. conducted experiments; U.D. and S.T. contributed to the proteomic analysis; O.B. and B.R. provided the SV2A KO mice; A.K and C.B. contributed to autophagy experiments; M.G., C.C., S.R. contributed to confocal and Super resolution microscopy, P.P. and B.P. conducted AlphoFold3 Modelling; M.J. P.P. and K.F. wrote the paper; K.F. supervised the project.

