## Supplementary Data for "The role of the synaptic vesicle protein SV2A in regulating mitochondrial morphology and autophagy"

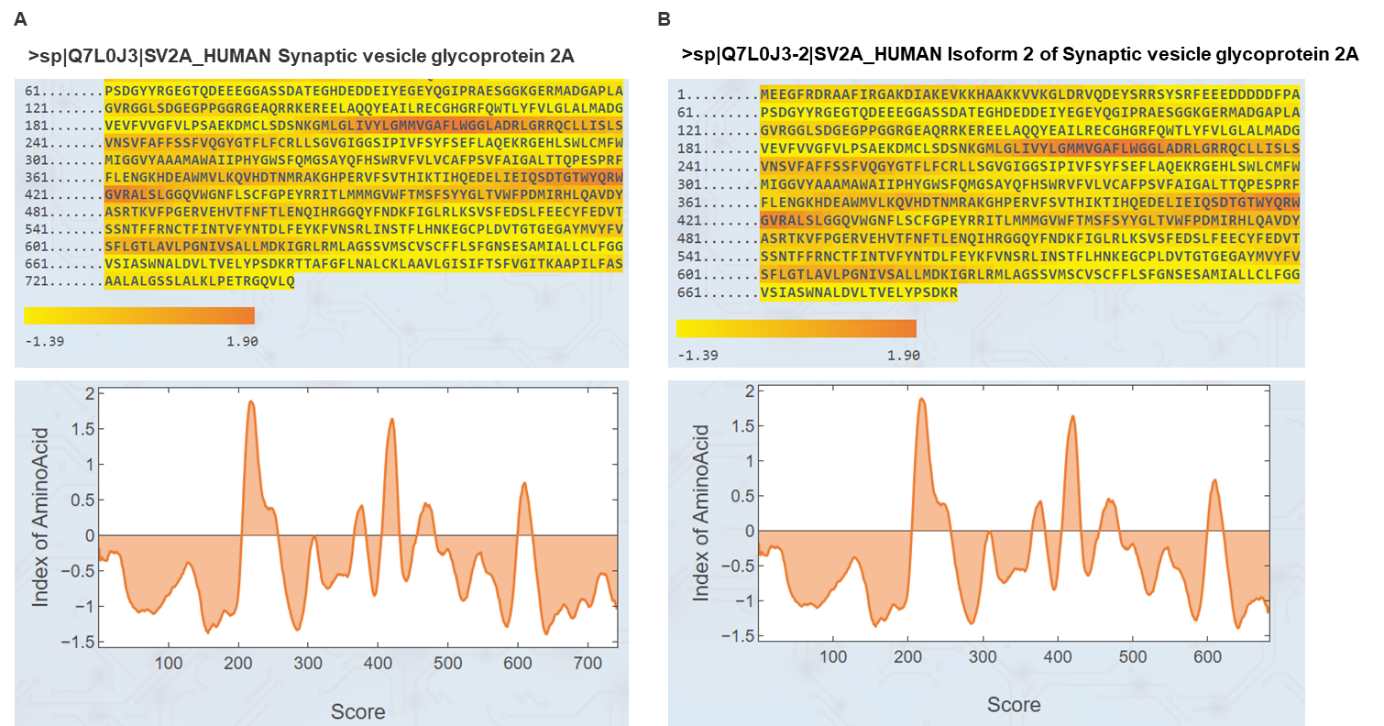

**Supplementary Figure 1 - The iMLP prediction shows the presence of an internal targeting signal in SVA and its isoform.**

A) iMTS-L propensity heatmap and its predicted iMTS-L propensity profile for SV2A and B) SV2A isoform.

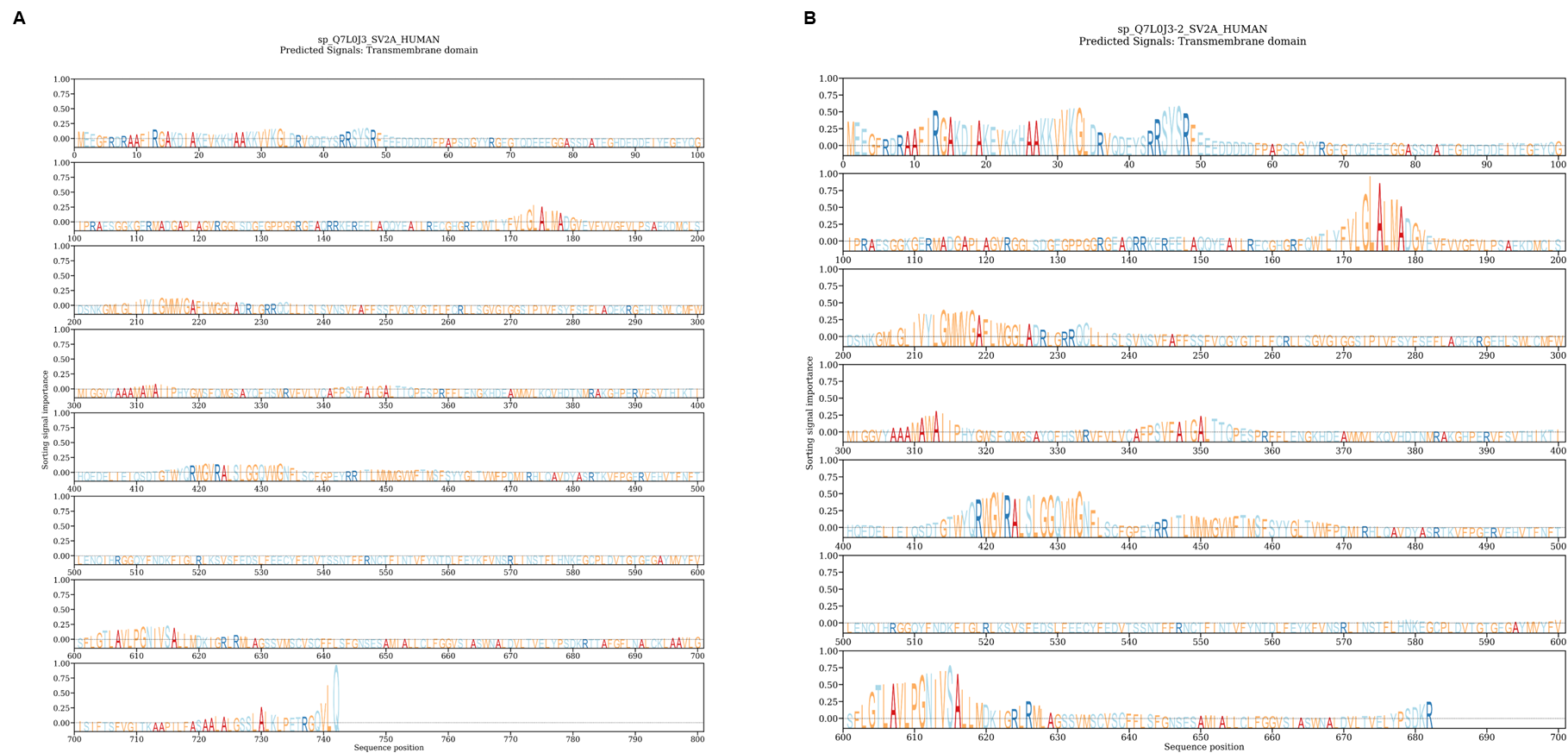

**Supplementary Figure 2 - Analysis of protein sequences of SV2A using DeepLoc2.0**.

A) The DeepLoc2.0 prediction of subcellular localization-associated sorting signals in SVA and its isoform (B).

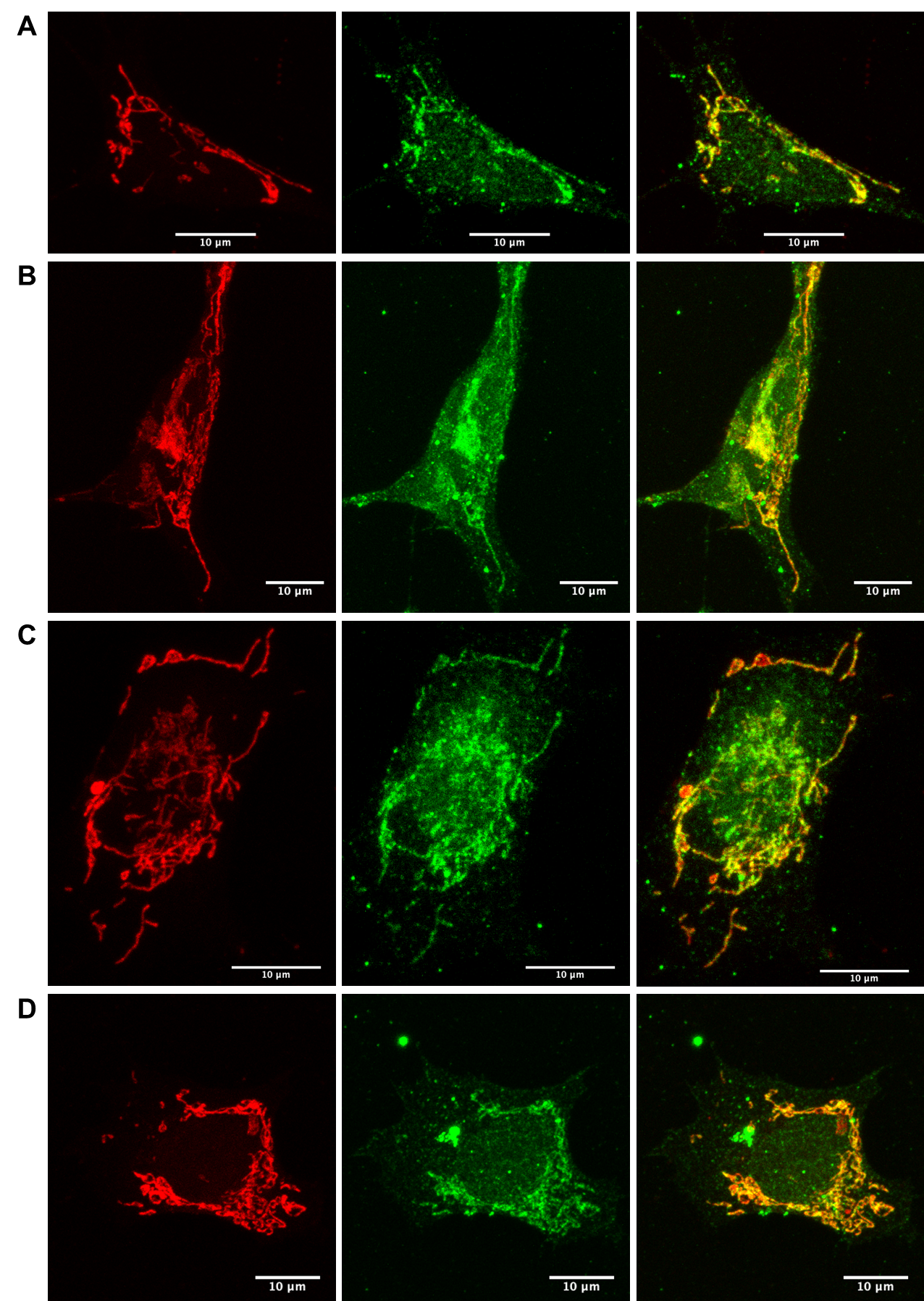

**Supplementary Figure 3 - SV2A is localized at the outer mitochondrial membrane.**

A-D Representative confocal fluorescence images of double-immunolabeled human SH-SY5Y cells stained for SV2A (green) and mitochondria (red) using MitoTracker™ Deep Red FM. Colocalization of SV2A and mitochondria is indicated by yellow spots in the merged image.

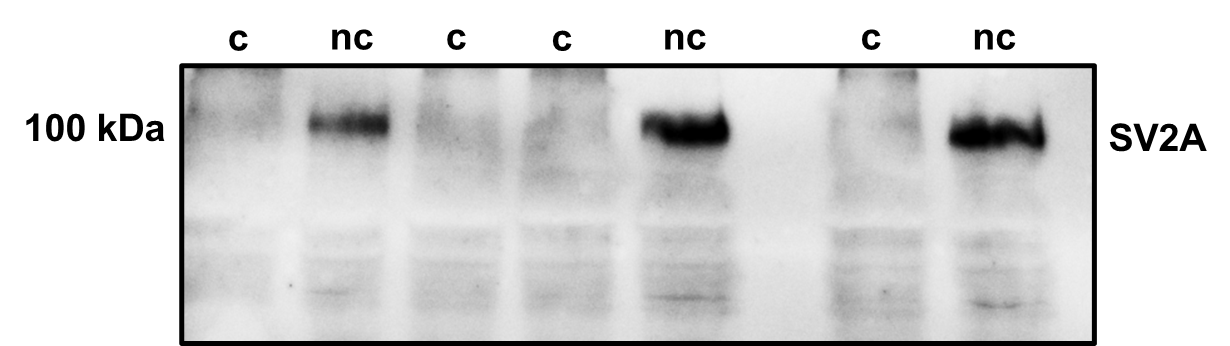

**Supplementary Figure 4 - Determination of Blotting conditions for SV2A.**

Uncooked (uc) and cooked (c) samples were compared using western blotting and transferred to PVDF membrane followed by SV2A antibody staining. Non-cooked band of SV2A are significant more present compared to cooked samples.

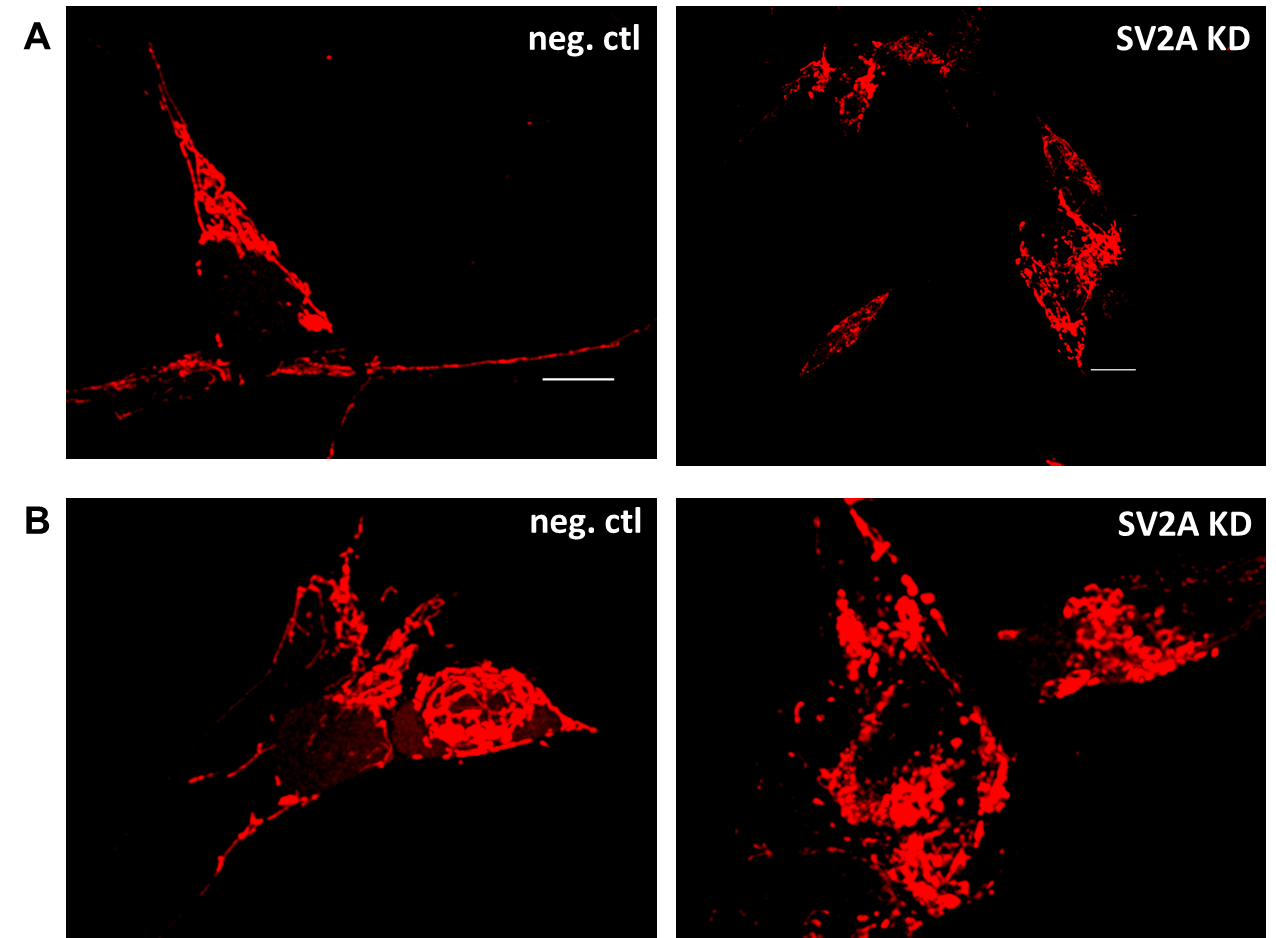

**Supplementary Figure 5 - Effect of SV2A KD on mitochondrial morphology.**

A-B Representative confocal images of mitochondria in siRNA-induced SV2A KD cells and cells treated with scrambled siRNA (10 nM/48 h). Length distribution of mitochondria in SV2A KD cells compared to cells treated with scrambled siRNA.

**Supplementary Table 1 - Analysis of interaction partners of SV2A using ReCLIP in HEK293 cells.**

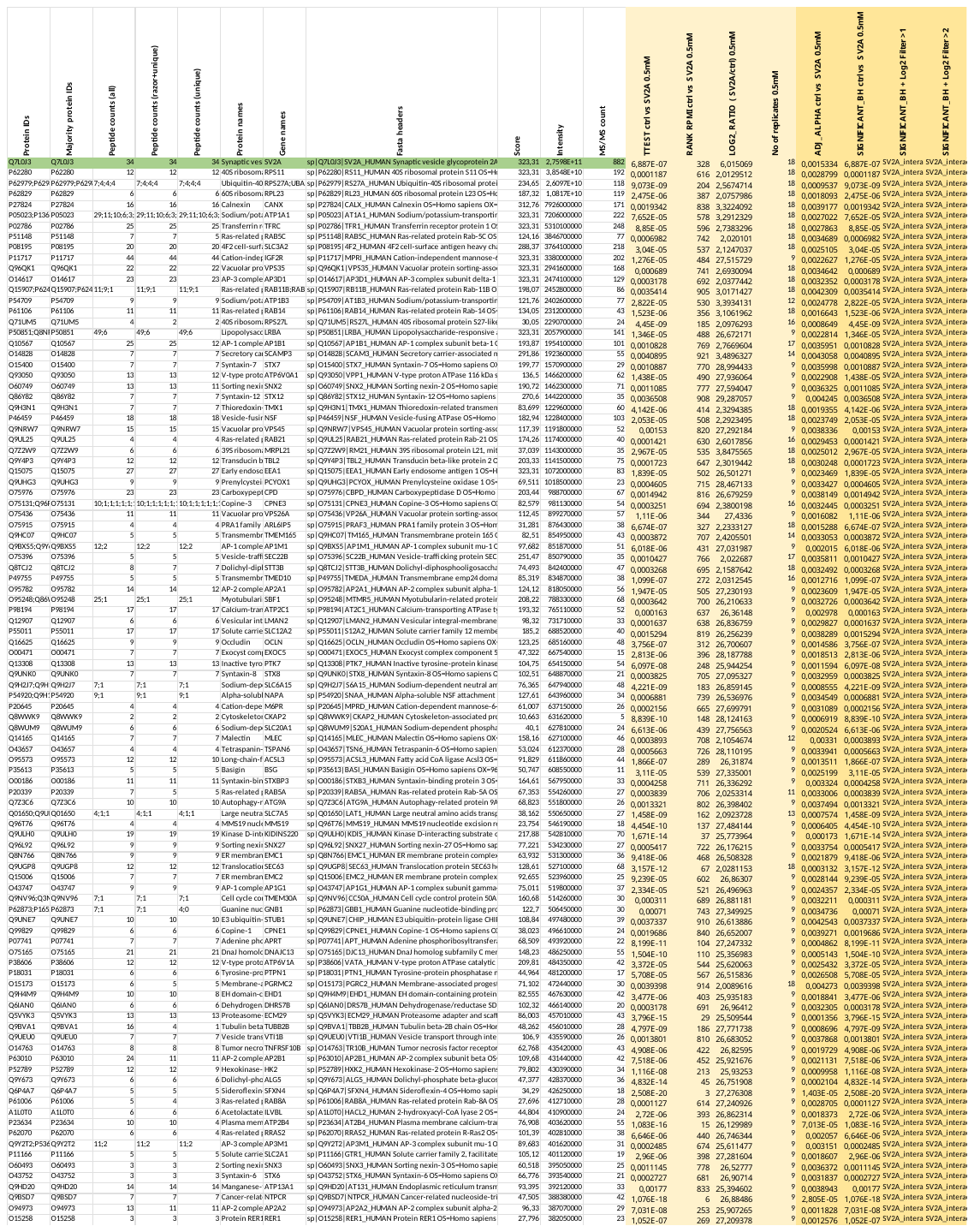

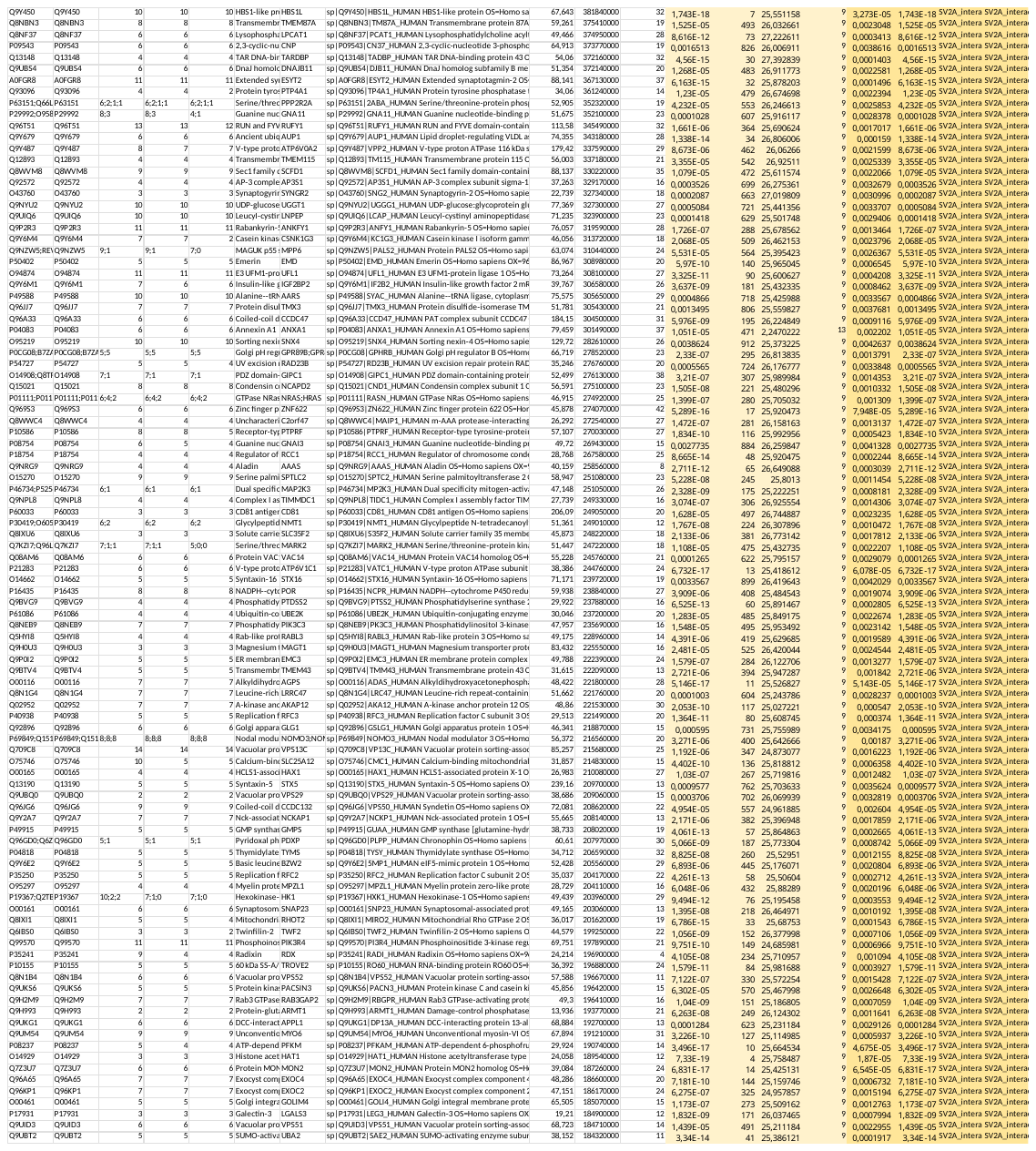

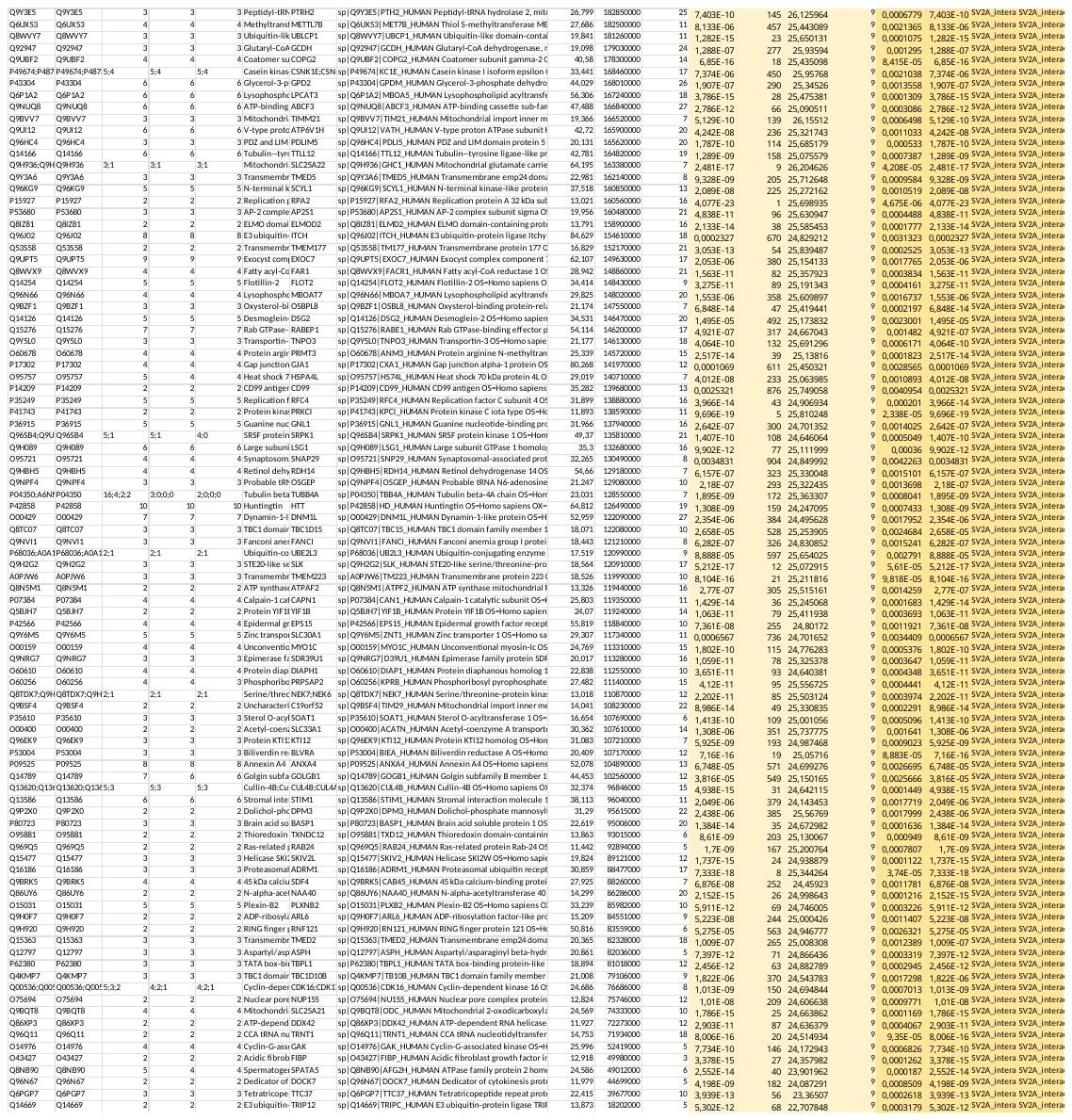
